# Chemical genetic screen for substrates of autism risk gene NUAK1 uncovers its role in ARF6 mediated dendritic spine maturation

**DOI:** 10.64898/2026.04.16.718295

**Authors:** Josilyn R. Sejd, Swagatika Paul, Daphnée M. Marciniak, Moira A. Cornell, Angel Sondhi, Shao-En Ong, Smita Yadav

## Abstract

Dysregulation of signaling by Novel (Nua) Kinase 1 (NUAK1) is associated with autism spectrum disorder, tissue fibrosis, and cancer progression. Direct phosphorylation targets of NUAK1 in the brain are unknown, hindering mechanistic understanding of its role in neurodevelopment. Here, we demonstrate that autism-associated NUAK1 variants differentially impact catalytic activity and subcellular distribution. We engineered ATP-analog sensitive NUAK1 and utilized its specificity towards bulky analogs to identify over 30 hitherto unknown direct phosphorylation targets of NUAK1 in the brain. We demonstrate that Pleckstrin Homology and Sec7-domain containing protein 3 (PSD3) is a *bona fide* phosphorylation target of NUAK1. PSD3, a guanine nucleotide exchange factor (GEF) for ARF6 GTPase, is phosphorylated by NUAK1 at Ser^476^. In neurons, expression of phosphodeficient PSD3 (S476A) leads to enhanced dendritic spine maturation in an ARF6-dependent fashion. Mechanistically, NUAK1 suppresses ARF6 activation through PSD3 phosphorylation. Abolishing Ser^476^ phosphorylation leads to aberrant ARF6 activation and increased generation of PI(4,5)P_2_. Our study reveals direct neuronal substrates of an autism risk gene NUAK1 and further delineates a mechanism through which NUAK1 phosphorylation of PSD3 controls ARF6 activation and dendritic spine maturation.

## Introduction

NUAK1 is a serine/threonine protein kinase that belongs to the AMP-activated protein kinase (AMPK) family(*1*). Despite the association of NUAK1 with pathophysiology of autism spectrum disorder (ASD) (*2*, *3*), neurodegenerative disorder (*4–6*), fibrosis (*7*, *8*), immune dysfunction (*9*), and cancer (*10–12*), our current understanding of NUAK1 signaling is limited. Several factors have hindered progress in elucidating the physiological function of NUAK1 in normal and disease states. *NUAK1* knockout mice are embryonically lethal (*13*), precluding investigation of its physiological function. Further, both overexpression and loss of function of NUAK1 are associated with pathogenesis suggesting the importance of its regulated activity. For example, high *NUAK1* expression is associated with reduced survival in pancreatic ductal adenocarcinoma and colorectal cancers (*9*, *12*). It is also upregulated in multiple fibrotic tissues in humans, and NUAK1 inhibition in mice attenuates fibrosis (*7*, *8*, *14*). NUAK1 stabilizes microtubule binding protein tau and in a tauopathy mouse model its deletion protects against neurodegeneration (*5*). In contrast, loss of NUAK1 in the brain causes defects in axonal development and heterozygous *NUAK1* knockout mice exhibit social behavior deficits (*15*, *16*). Physiological direct phosphorylation targets of NUAK1 have not been identified, with a few exceptions, constraining understanding of NUAK1 function in normal and disease states.

Primarily studied in context of tumor suppression and cell proliferation (*10*, *17–19*), NUAK1 contains 661 amino acids with an N-terminal kinase domain and a largely disordered C-terminal region. NUAK1 activity is regulated by LKB1 (liver kinase B1) and NDR2 (Nuclear df2-related kinase 2) phosphorylation at residue Thr^211^ in its activation loop (*20*, *21*). Phosphorylation by AKT (Protein kinase B) at Ser^600^ has been shown to further increase the catalytic activity of NUAK1 (*22*). The LKB1-NUAK1 mediated signaling controls cell adhesion through regulation of myosin phosphatase complexes (*17*). Three Gly-Iso-Leu-Lys (GILK) motifs in NUAK1 mediate its interaction with the phosphatase complex MYPT1-PP1β (Myosin phosphatase targeting subunit 1/Protein phosphatase 1 beta) that controls cell adhesion through regulation of myosin light chain (*17*). An alternative pathway for NUAK1 phosphoregulation mediated by calcium-dependent activation of PKCα has been implicated in cancer cells lacking LKB1 (*23*, *24*), highlighting several routes for achieving NUAK1 activation in distinct cellular context. Greater complexity to NUAK1 signaling arises from distinct subcellular compartments in which NUAK1 has been reported to function. A bipartite nuclear localization signal shuttles NUAK1 into the nucleus, with oxidative stress increasing its nuclear transport (*25*). In cancer cell lines, NUAK1 appears to be predominantly nuclear, localized within nuclear speckles where it plays an important role in regulating splicing (*24*). Therefore, regulation of NUAK1 signaling and its phosphorylation substrates depend on cellular context.

While essential for normal brain development, NUAK1 neuronal function and signaling is less understood. Highly expressed in the central nervous system, NUAK1 has been shown to be important for axonal and dendrite development (*15*, *16*, *26*, *27*). Catalytic activity of NUAK1 is essential for its function in the neuronal development. For example, heterozygous *NUAK1* knockout mice exhibit impaired axon elongation which can be rescued by expression of active NUAK1, but not by kinase-dead NUAK1 (*16*). Further, small molecule kinase inhibitors against NUAK1 impair both axon and dendritic development (*26*). However, mechanisms through which NUAK1-mediated phosphorylation promotes neuronal development are unknown largely because its direct phosphorylation targets in the brain have not been identified. Further, whether ASD associated NUAK1 variants impact its catalytic activity have not been tested.

In our study, we biochemically characterized ASD-associated NUAK1 variants and show their differential impact on catalytic function and subcellular localization. Given the importance of its kinase activity and dearth of mapped NUAK1 neuronal substrates, we employed an unbiased chemical-genetic strategy to identify direct phosphorylation targets of NUAK1 in the mouse brain. We reveal neuronal substrates of NUAK1 kinase that span cellular processes of endocytosis, splicing, axon development, dendritic growth, and synaptogenesis. From these candidate substrate hits, we demonstrate that PSD3 (Pleckstrin Homology and Sec7-domain containing protein 3), a guanine exchange factor (GEF) for the small GTPase ARF6 (ADP-ribosylation factor 6), is a *bona fide* substrate of NUAK1. Through a combination of in vitro biochemical assays, phosphomutant analyses in neurons, and superresolution imaging, we have uncovered an essential role of NUAK1 mediated phosphorylation of PSD3 that controls ARF6 and PI(4,5)P_2_ dependent dendritic spine maturation.

## Results

### Autism-Associated NUAK1 Variants Perturb Kinase Activity and Subcellular Distribution

NUAK1 has been reported to have differential subcellular localization to the nucleus and/or cytosol depending on cell type and nutrient conditions. To study the role of NUAK1 in brain development, we first investigated its localization in neuronal cells. Rat embryonic hippocampal neurons were fixed at 14 days in vitro (DIV) and then immunostained with an antibody against NUAK1 to visualize localization of endogenous NUAK1, along with an antibody against the dendrite marker MAP2 (microtubule associated protein 2) (Fig.1A). We found that NUAK1 is localized to the cytosol within the soma, dendrites, and axon within neurons (Fig.1A). In human brain, the Human Brain Transcriptome database (*28*) shows *NUAK1* mRNA is also highly expressed in early development across all measured brain regions including the neocortex, and its expression persists into adulthood (Fig.1B). Similarly, *NUAK1* mRNA is highly expressed in neurons within the cortex, hippocampus, and cerebellum as revealed by in situ hybridization in the Allen Brain Atlas database (Fig.1C).

**Figure 1.**
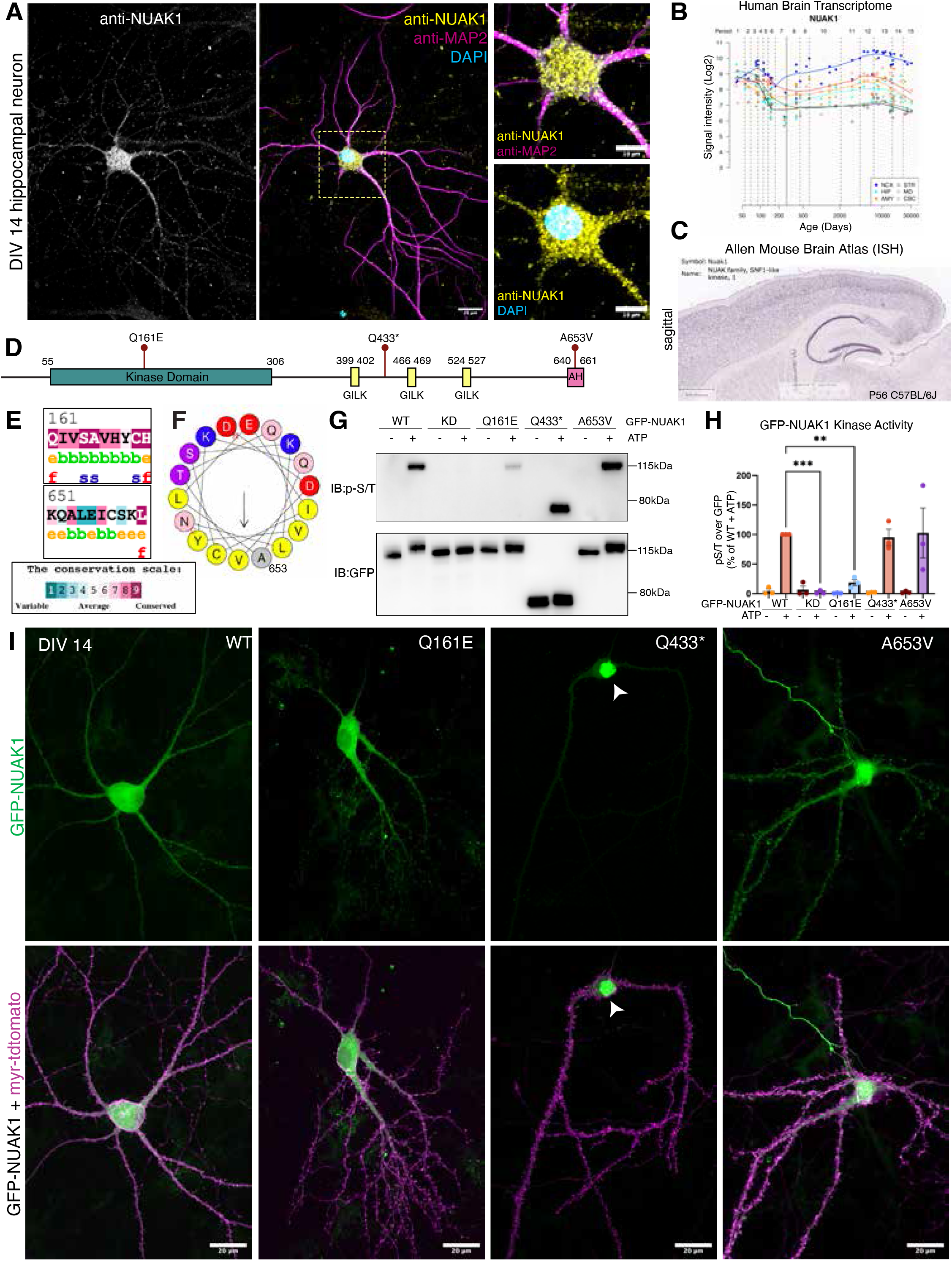
Autism-associated NUAK1 mutations impact catalytic activity and subcellular localization. **(A)** Day in vitro (div) 14 rat hippocampal neurons stained with antibodies against NUAK1 and MAP2, with nuclear marker DAPI. Right panels show magnified soma with anti-NUAK1 and either MAP2 (top panel) or DAPI (bottom panel). Error bars represent 20µm in full images and 10µm in zoomed images at right. **(B)** Human brain transcriptomic data for NUAK1 obtained from Human Brain Transcriptome (HBT) database across different stages of human brain development. Genome-wide, exon-level transcriptome data in HBT was generated using the Affymetrix GeneChip Human Exon 1.0 ST Arrays from over 1,340 tissue samples sampled from both hemispheres of postmortem human brains. Specimens range from embryonic development to adulthood (periods 1-15) and represent both males and females. Brain regions plotted are the neocortex (NCX), hippocampus (HIP), amygdala (AMY), striatum (STR), mediodorsal nucleus of the thalamus (MD), and cerebellar cortex (CBC). **(C)** In situ hybridization data from the Allen Brain Atlas using antisense probe against NUAK1 on sagittal sections of postnatal day 56 (P56) C57BL6/J male mouse brain. Scale bar represents 839µm. **(D)** Two-dimensional schematic of NUAK1 protein domains, with the N-terminal kinase domain, 3 GILK motifs, and C-terminal amphipathic helix indicated with colored boxes. The three NUAK1 ASD-associated mutations are shown with lollipops above the protein domains: the two missense mutations Q161E and A653V, and the early truncation mutation Q433ter (Q433*). **(E)** ConSurf evolutionary conservation analysis of the two ASD-associated missense mutations in NUAK1: Q161E and A653V. Residues are colored according to their evolutionary conservation, with warm colors indicating conserved residues and cool colors indicating variable residues. **(F)** Heliquest-predicted amphipathic helix at the C-terminus of NUAK1, which spans amino acids 644-661. Ala^653^, the site of the A653V ASD-associated mutation, is indicated. The arrow indicates the hydrophobic face of the amphipathic helix. **(G)** Western blot of immunoprecipitated GFP-NUAK1 mutants incubated plus/minus ATP to allow NUAK1 to autophosphorylate. Immunoblots show anti-phospho-serine/threonine and anti-GFP (loading control). **(H)** Quantification of kinase activity of NUAK1 mutants, with the phospho-serine/threonine signal divided by the GFP loading control and all samples normalized as a percentage of WT + ATP per experiment (n=3 from three biological replicates, ** denotes p<0.005; *** denotes p<0.001, one way ANOVA with Tukey’s multiple comparisons). All error bars represent standard error of the mean. **(I)** DIV14 rat embryonic hippocampal neurons expressing GFP-tagged NUAK1 mutants and membrane marker myristoyl-tdtomato (myr-tdtomato). Neurons were transfected with GFP-NUAK1 and myr-tdTomato on DIV12 and fixed 48 hours later. The arrowhead in Q433* image indicates the predominantly nuclear localization of this mutation. Scale bar represents 20μm in all images.

We next interrogated the impact of ASD-associated variants on the catalytic activity and neuronal localization of NUAK1 kinase. Three NUAK1 variants have been reported to date: missense mutation Q161E within the kinase domain, premature truncation mutant Q433*, and missense mutation A653V in the C-terminal tail (*29–31*) (Fig.1D). Evolutionary analyses with ConSurf (*32*) revealed conservation of the residues mutated in missense mutations. The Gln^161^ residue shows 100% conservation across 150 species, suggesting that this residue is important for NUAK1 activity and that the mutation to the charged glutamate residue will affect NUAK1 activity. The Ala^653^ residue shows 98% conservation, with only the small amino acids glycine or serine observed at this position in other species (Fig.1E). Alphafold3 (*33*, *34*) model of NUAK1 predicts Ala^653^ residue to be located in an alpha helix. We used HeliQuest (*35*) to probe the nature of this alpha helix and found that residues 640-661 formed an amphipathic helix (Fig.S1A-B), with Ala^653^ located in the hydrophobic face (Fig.1D and 1F). The mutation of Ala^653^ to Val may interfere with NUAK1 binding with other proteins or its subcellular localization.

Next, we tested whether autism-associated NUAK1 variants impacted catalytic activity by performing an in vitro kinase activity assay (*36*). Briefly, GFP-tagged NUAK1 variants were expressed in HEK293T cells, followed by lysis, immunoprecipitation, and incubation with Mg^2+^ and ATP. We observed robust autophosphorylation with wild-type (WT) NUAK1, which was lost with the catalytic lysine K84M kinase dead (KD) mutation (Fig.1G-H). We observed a greater than 80 percent reduction in catalytic activity with the Q161E mutant, while the Q433* and A653V mutations had unchanged kinase activity relative to WT NUAK1 (Fig.1G-H). To evaluate the impact of NUAK1 variants on its subcellular distribution in neurons, we expressed GFP-tagged NUAK1 mutants in hippocampal neurons. Similar to endogenous NUAK1 protein, exogenously expressed green fluorescent protein (GFP)-tagged NUAK1 was also largely cytosolic (Fig.1I). We found that while the Q161E and A653V variants had similar localization to WT, expression of NUAK1 truncation variant Q433* led to over four-fold enrichment to the nucleus over cytoplasm (Fig.1I and Fig.S1D). Because the A653V mutation did not impact catalytic activity or nuclear localization in neurons, we sought to ascertain if any other functions of this protein were impacted by this mutation. NUAK1 in mitotically active cells has well-established kinase activity-dependent roles in the regulation of RNA splicing (*24*, *37*). WT NUAK1 has been shown to colocalize with SC35 (SRSF2 or Serine/arginine-rich splicing factor 2), a marker of spliceosomes, in the nucleus in various contexts (*24*, *37*). We expressed GFP-tagged NUAK1 (WT, Q161E, Q433* and A653V) in human WTC11 (Corielle) iPSC-derived neural progenitor cells (NPCs) (Fig.S1C). In NPCs, WT NUAK1 localizes to the cytosol, nucleus and is also enriched in nuclear speckles. In contrast, truncation mutant Q433* NUAK1 is entirely absent from the cytosol and instead enriched in nuclear speckles positive for SC35. Remarkably, the kinase deficient mutant Q161E NUAK1 does not show colocalization with SC35 positive nuclear speckles. While catalytically active, in NPCs, the mutant A653V NUAK1 shows nuclear and cytosolic localization but is absent from SC35 positive nuclear speckles (Fig.S1C). These experiments suggest that NUAK1 mutations might lead to decreased phosphorylation of its neuronal substrates either through deficits in its catalytic function (Q161E) or aberrant shift in its distribution between the cytoplasm to the nucleus/nuclear speckles (A653V and Q433*).

### Chemical-Genetic Screen for Identification of NUAK1 direct neuronal targets

NUAK1 phosphorylation targets in the developing brain have not been identified. We employed a chemical-genetic approach (*38–41*) where we engineered a bulky ATPγS analog sensitive (AS) NUAK1 that uniquely thiophosphorylates substrates and can be covalently captured by iodoacetamide beads for identification by mass spectrometry. The advantages of this method are that first, it identifies direct phosphorylation targets of the kinase of interest, and second, the phosphorylation site on the substrate that is revealed can be then modified to study the physiological impact of phosphorylation. To engineer the AS-NUAK1 kinase, we mutated the conserved gatekeeper residue Met^132^ in the kinase domain of NUAK1 (Fig.2A) to a smaller amino acid Ala or Gly. With the enlarged hydrophobic nucleotide binding pocket, the engineered AS-NUAK1 can utilize N^6^-substituted bulky ATP S molecules and thiophosphorylate its substrates. As a negative control, we also generated kinase dead (KD) NUAK1 with a Lys84Met (K84M) mutation, abolishing the conserved catalytic lysine residue that is responsible for coordinating the - phosphate of ATP. We bacterially purified the kinase domain of NUAK1 (residues 1-433) with either active wildtype (WT), kinase-dead (K84M), and analog sensitive (M132G) mutations (Fig.S2A-C). We next tested different NUAK1 AS mutants and bulky ATP analogs combinations, and determined that N^6^-furfuryl-ATP S is optimally utilized by M132G NUAK1 for phosphorylation but not by WT NUAK1 (Fig.2B-C). We confirmed that full-length NUAK1 M132G shows preferential utilization of N^6^-furfuryl-ATP S using full-length HA (hemagglutinin)-NUAK1 immunoprecipitated from HEK293T cells (Fig.S2D-E).

**Figure 2.**
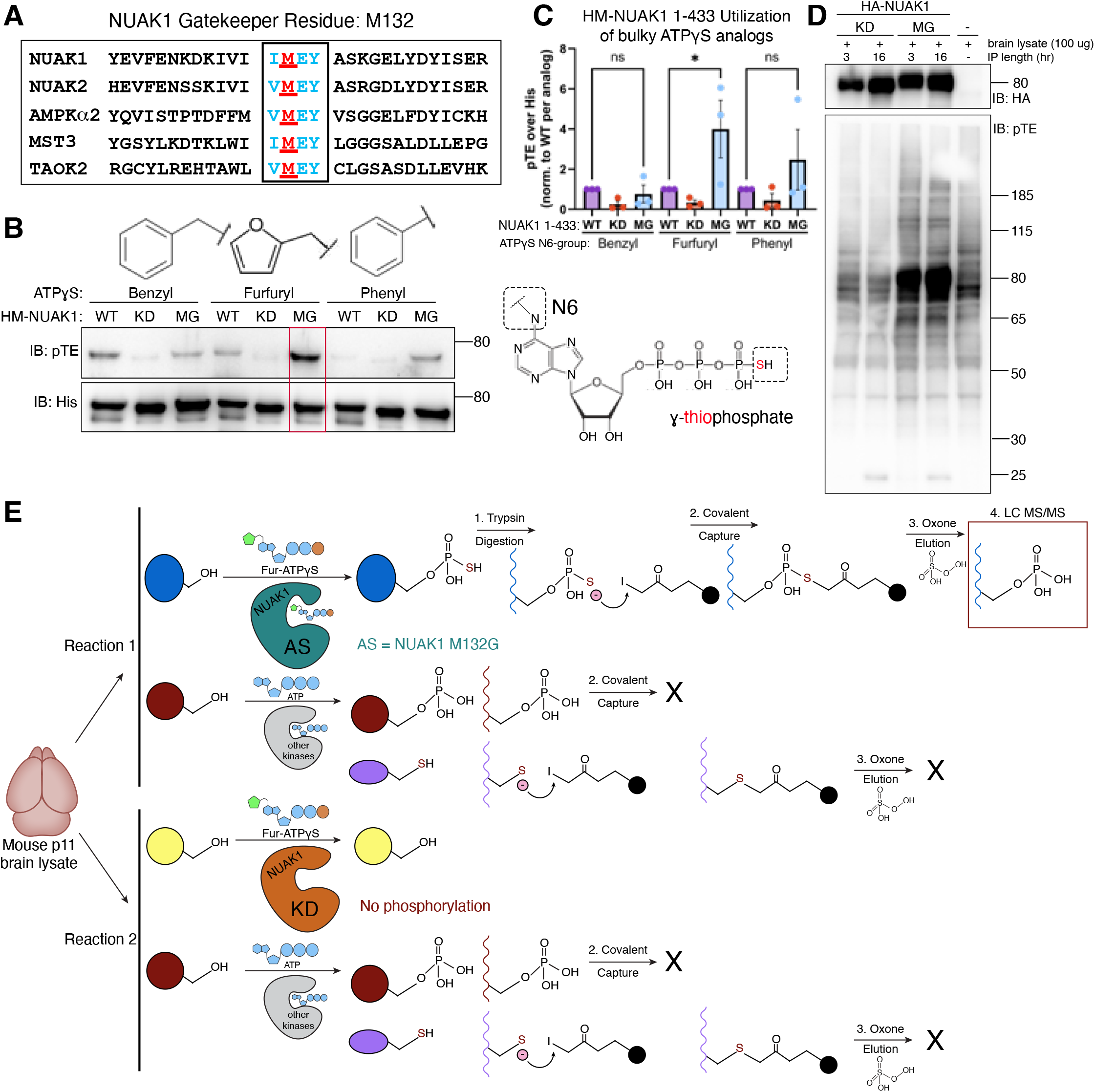
Engineering ATP-analog sensitive NUAK1 kinase through the gatekeeper residue mutation M132G. **(A)** Multiple sequence alignment of the kinase domains of serine/threonine kinases, including NUAK1, NUAK1, AMPK 2, MST3, and TAOK2. The conserved methionine gatekeeper residue is colored red and underlined, while the flanking residues are colored blue. The four highly conserved residues surrounding the gatekeeper are indicated with a box. **(B)** Top: bulky side groups (benzyl, phenyl, and furfuryl) that are N6-substitutents of ATP S (right) in kinase assays. Bottom: Western blots showing utilization of bulky ATP S molecules by purified HM-tagged NUAK1 (amino acid 1-433). Antibody against phosphothioester (pTE) is used as a readout of autophosphorylation, while anti-His is used as a loading control. The optimal gatekeeper mutant/bulky ATP S combination of M132G NUAK1 and furfuryl-ATP S is indicated with a red box. **(C)** Quantification of utilization of bulky ATP S analogs by HM-NUAK1, with pTE signal divided by loading control His for each sample. Data were normalized to WT for each analog, with KD and MG represented as a ratio of WT (n=3 biological replicates, one-way ANOVA with Bonferroni’s multiple comparison test, * denotes p<0.05). Error bars represent standard error of the mean. **(D)** HA-tagged full-length NUAK1 KD or MG was expressed in HEK293T cells followed by lysis and immunoprecipitation for the indicated time. Immunoprecipitated NUAK1 was incubated with mouse brain lysate and furfuryl ATP S, followed by alkylation to generate phosphothioester for WB detection. HA is used as a loading control for NUAK1, while phosphothioester (pTE) is used as a readout of substrate phosphorylation in mouse brain lysate. **(E)** Schematic of labeling of mouse brain lysate by analog-sensitive (AS) NUAK1 and N6-furfuryl-ATP S (reaction 1), or negative control K84M (KD) NUAK1 and N6-furfuryl-ATP S (reaction 2). Thiophosphorylated substrate proteins were digested with trypsin, and thiophosphorylated peptides were then covalently captured using iodoacetamide (IAA) beads. Beads were treated with oxone to specifically elute off thiophosphorylated peptides as phosphopeptides, while peptides with other thiol groups (such as cysteines) are retained on the beads. The resulting phosphopeptides are then analyzed by LC-MS/MS.

Since motifs outside the NUAK1 kinase domain such as the “Gly-Iso-Leu-Lys” (GILK) motif are important for substrate binding, we utilized full-length AS-NUAK1 (M132G) to identify thiophosphorylated substrates from mouse brain lysate. We affinity purified full-length AS-NUAK1(M132G) and kinase-dead NUAK1(K84M) from HEK293T cells using HA-conjugated beads and incubated them with mouse brain lysate in the presence of N^6^-furfuryl ATP S (Fig.2D). Compared to the lysate only or kinase dead controls, we observed robust signal for thiophosphorylated substrates identified by a phosphothioester antibody in the AS samples but not in KD or lysate only samples. After validating that full-length HA-NUAK1 AS is able to effectively phosphorylate substrates using N^6^-furfuryl ATP S, we scaled up our system for mass spectrometry analysis. Immunoprecipitated NUAK1 was incubated with mouse brain lysate and N^6^-furfuryl ATP S to allow AS-NUAK1 to thiophosphorylate its substrates. Mouse brain lysate was obtained from pooled tissue from 8 individual p13-15 mouse pups. Our experimental design consisted of 5 kinase dead NUAK1 samples and 5 analog-sensitive NUAK1 samples, each incubated with mouse brain lysate and furfuryl ATP S. This resulted in 5 biological replicates for each condition. Samples were then denatured with urea and digested to generate peptides. The resulting thiopeptides were then captured on iodoacetamide resin and treated with oxone to specifically elute peptides containing a thiophosphate moiety, while cysteine (thiol) containing peptides were retained on beads. Eluted phosphopeptides were separated using liquid chromatography and analyzed using mass spectrometry (Fig.2E). Using this workflow, our mass spectrometry analyses of five replicate samples revealed 972 total phosphopeptides.

### Identification of NUAK1 Consensus Sequence and Direct Neuronal Substrates

Our criteria for narrowing candidate hits from the chemical genetic screen was that the substrate should be present in at least three out of five AS replicates and also be absent in (0/5) all kinase dead (KD) replicates. From our list of 972 total phospho-serine and phospho-threonine peptides, 31 peptides matched this stringent criterion. These 31 potential NUAK1 substrate proteins were analyzed for their cellular function and were binned into the following categories: endocytic recycling, splicing, cytoskeletal reorganization, as well as axonal, dendritic, and synaptic development (Fig.3A). Proteins were assigned to one or more categories based on their previously established roles in each cellular process and color coded with either a single color or a gradient of 2-4 colors to indicate their pleiotropic functions (Fig.3A). We used the sequence of these 31 peptides to generate a sequence logo using Phosphosite Plus to show the preferred phosphorylation motif of NUAK1 in our mass spectrometry experiment (Fig.3B). We further shortlisted peptides where the phosphosite sequence fit the NUAK1 consensus sequence determined previously through peptide arrays (*42*). Out of the 31 phosphopeptides, 17 had a basic residue at the −3 position (Fig.3C). Our data demonstrate that the stringent consensus sequence for NUAK1 is -[R/K/H]-[R/K/H]-S-[R/K/H]-pS*-X-X-X-where is an aliphatic residue (Fig.3C). The function and subcellular localization of the 17 putative NUAK1 substrates are listed and both nuclear and cytosolic proteins were identified in the chemical genetic screen (Fig.3C).

**Figure 3.**
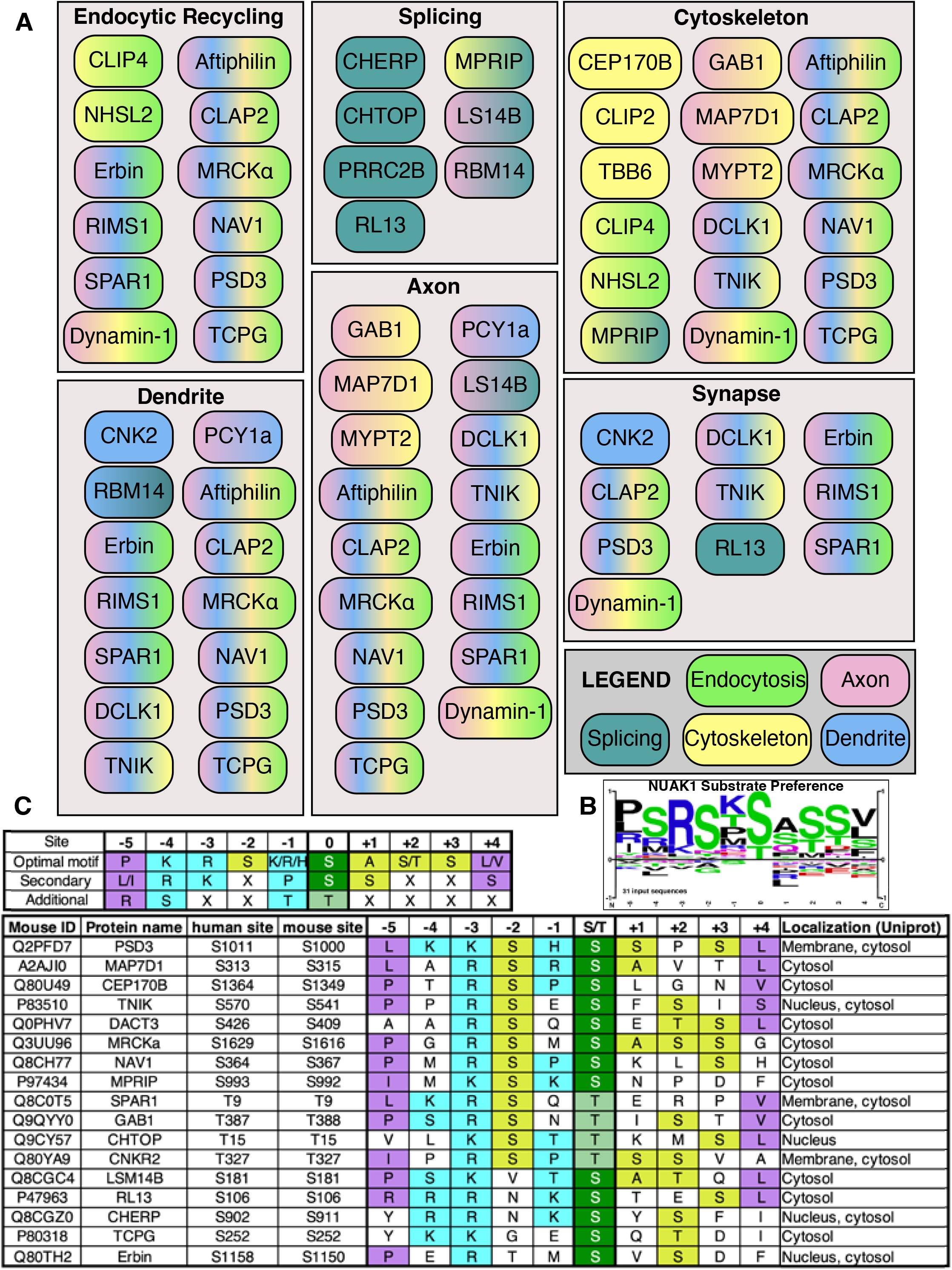
NUAK1 phosphorylation targets identified by chemical genetic screen. **(A)** Putative NUAK1 substrate proteins (31 total) that occurred in at least 3/5 analog sensitive samples and 0/5 kinase dead samples were analyzed for their roles in endocytosis, splicing, cytoskeletal reorganization, as well as development of dendrites, axons, or synapses. Each substrate is color-coded according to its role(s) in each of these processes, with light green for endocytosis, teal for splicing, yellow for cytoskeleton, pink for axon, and blue for dendrites, with the legend in the lower right panel showing each individual color. Substrates with roles in synaptic development are located in the “synapse” box on the middle right. Substrates with multiple functions are indicated with gradients with 2-4 colors. **(B)** The 9 amino acids surrounding the phosphoacceptor (from residues 5 to +4) for the 31 potential substrates of NUAK1 that occurred in at least 3/5 analog sensitive samples and 0/5 kinase dead samples were analyzed using the Phosphosite.org WebLogo tool to generate a preference map for NUAK1 substrate phosphorylation. **(C)** Putative NUAK1 substrate proteins that occurred in at least 3/5 analog sensitive samples and 0/5 kinase dead samples were analyzed for how well they fit the established NUAK1 consensus sequence. Most stringently, NUAK1 requires a basic residue (K/R) at the −3 position. After filtering the 31 unique potential substrates by this stringent consensus sequence, 17 putative substrates remained. The mouse Uniprot ID and protein name for each protein is listed, as are the human and mouse phosphosites. The amino acids surrounding the phosphosite are listed in this table, from the −5 position to the +4 position relative to the phosphoacceptor. Based on the established NUAK1 consensus sequence, along with the data from our MS runs, we were able to establish the optimal motif for substrate phosphorylation by NUAK1. This optimal motif: (1) contains a basic residue at −3 position; (2) contains a serine at −2 position; (3) contains a serine phosphoacceptor; and (4) contains an aliphatic branched chain amino acid at −5 and +4 positions. Residues are color coded based on whether the residue fits the optimal NUAK1 motif at the given site, with positions −5 and +4 in purple, positions −4, −3, and −1 in cyan, and positions −2, +1, +2, and +3 in chartreuse. The central phosphoacceptor at position 0 is colored in dark green for serine (more preferred) and light green for threonine (less preferred).

### PSD3 is a Bona Fide Substrate of NUAK1 Kinase

We next focused on our top candidate in the screen for NUAK1 substrates, Pleckstrin Homology Domain and Sec7 domain-containing protein 3 (PSD3). A multidomain protein, PSD3 consists of a pleckstrin homology (PH) domain which binds specifically to phospholipid PI(4,5)P_2,_ and a sec7 domain that acts as a guanine exchange factor (GEF) for the small GTPase ARF6 (*43*, *44*). A coiled coil (CC) region following the PH domain enhances membrane association of the PH domain and has been shown to bind actin (*45*, *46*). These 3 functional domains are present in the alternatively spliced isoforms of human PSD3 (Fig.4A). The canonical isoform 1 contains 1048 amino acids (Uniprot #Q9NYI0), while isoform 2 has a single amino acid deletion and contains 1047 amino acids (Uniprot #Q9NYI0-2). Isoform 3 is 513 amino acids long but contains the conserved Sec7 domain, PH domain and CC region (Uniprot #Q9NYI0-3). The putative NUAK1 phosphorylation site Ser^1011^ is present following the CC region and is shared by all three isoforms (Fig.4A). Previous studies have shown that the predominant isoform of PSD3 expressed in the mouse brain is 56kD, which corresponds to isoform 3 (*45*, *47*). We utilized ConSurf (*32*) to perform evolutionary conservation analysis, which revealed that the C-terminus of PSD3 is highly conserved, specifically around the PH, Sec7, and CC domains. Further, the phosphosite Ser^476^ and the motif surrounding it are extremely conserved (Fig.4B and Fig.S3A). Because our MS-identified phosphosite is located within the evolutionarily conserved C-terminus of both isoforms, we elected to utilize the more abundant shorter isoform 3 for further analyses.

**Figure 4.**
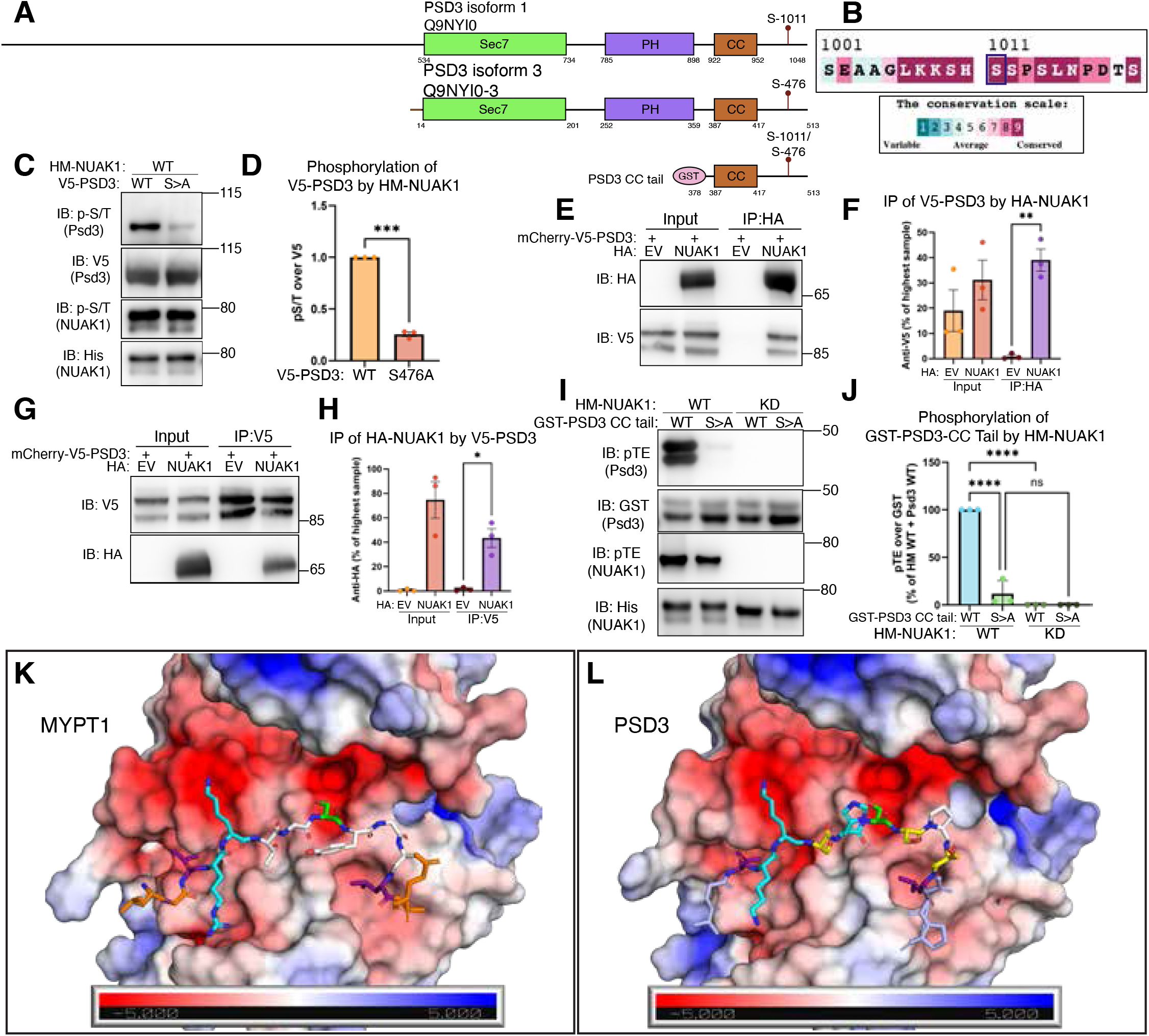
PSD3 is a bona fide direct phosphorylation target of NUAK1 kinase. **(A)** Two-dimensional schematic representation of the function domains of PSD3 isoform 1 (Uniprot # Q9NYI0), isoform 3 (Uniprot # Q9NYI0-3), and the shortened PSD3 construct used for biochemical experiments, PSD3 CC tail. PSD3 isoforms 1 and 3 share the same 3 conserved functional domains, including a Sec7 GEF domain (green) which activates ARF6, Pleckstrin Homology (PH) domain with specificity for PI(4,5)P_2_, and a Coiled Coil (CC) domain which binds to actin. The MS-identified phosphosite on PSD3, Ser^1011^ in isoform 1 or Ser^476^ in isoform 3, is indicated with a lollipop. The shortened CC tail domain used for protein purification contains an N-terminal GST tag followed by the C-terminal region of PSD3, from the CC domain to the end of the protein containing the Ser^476^/Ser^1011^ phosphosite. **(B)** ConSurf evolutionary conservation analysis of the MS-identified phosphosite of PSD3, Ser^1011^ in isoform 1. The Ser^1011^ phosphoacceptor residue is boxed in violet. Residues are colored according to their evolutionary conservation, with warmer colors indicating conserved residues and cool colors indicating variable residues. **(C)** V5-tagged PSD3 isoform 3 WT or S476A was immunoprecipitated from HEK293T cells, followed by incubation with purified HM-NUAK1 WT and ATP. Immunoblots show V5 as a loading control for PSD3 and His as a control for NUAK1, and phospho-serine/threonine as a readout of phosphorylation for both NUAK1 and PSD3 at their respective molecular weights. **(D)** Quantification of phosphorylation of PSD3 by NUAK1, as measured by phospho-serine/threonine divided by V5 signal. Each experiment is normalized to WT PSD3, with S>A PSD3 as a ratio of WT signal (n=3 biological replicates; *** denotes p<0.001, student’s t-test). Error bars represent standard error of the mean. **(E)** HEK293T cells were co-transfected with mCherry-V5-PSD3 isoform 3 and either HA-empty vector or HA-NUAK1 WT. Cells were then lysed, and protein was immunoprecipitated using HA beads. Immunoblots show HA signal for HA-empty vector or HA-NUAK1, and V5 signal for mCherry-V5-PSD3. **(F)** Quantification of V5 signal for input and HA IP for N=3 independent biological replicates. Error bars represent standard error of the mean; ** denotes p<0.005, one way ANOVA with Tukey’s multiple comparisons. **(G)** HEK293T cells were co-transfected with mCherry-V5-PSD3 isoform 3 and either HA-empty vector or HA-NUAK1 WT. Cells were then lysed, and protein was immunoprecipitated using V5 beads. Immunoblots show HA signal for HA-empty vector or HA-NUAK1, and V5 signal for mCherry-V5-PSD3. **(H)** Quantification of HA signal for input and V5 IP, for N=3 independent biological replicates. Error bars represent standard error of the mean; * denotes p<0.05, one way ANOVA with Tukey’s multiple comparisons. **(I)** HM-NUAK1 WT or KD was incubated with GST-PSD3-CC tail WT or S>A in the presence of Mg^2+^ and ATP S, followed by alkylation with PNBM to generate thiophosphate esters as a readout of thiophosphorylation by NUAK1. **(J)** Quantification of phosphorylation of GST-PSD3-CC tail WT or S>A by HM-NUAK1 WT or KD. Data are normalized to WT PSD3 + WT HM-NUAK1 for each experiment, with all other samples as a percentage of WT/WT (n=3 independent biological replicates; **** indicates p<0.0001, one way ANOVA with Tukey’s multiple comparisons). Error bars represent standard error of the mean **(K)** Alphafold3-predicted structure of NUAK1 WT (Uniprot # O60285) with the previously identified and validated substrate MYPT1 (Uniprot ID #O14974), along with requisite adaptor subunit Protein Phosphatase 1 beta (Uniprot ID #P62140). NUAK1 kinase domain is displayed as an electrostatic surface representation, with red-toned colors representing a negative charge, and blue-toned colors representing a positive charge. MYPT1 is colored in light blue, and amino acids within the NUAK1 substrate binding groove are color coded according to their position and side chain, with Leu at −5 and +4 in limon, Lys/His at −4, −3, and −1 in blue, and central phospho-acceptor Ser in green. Amino acid side chains are shown for positions −5 to +4 surrounding the MYPT1 Ser^445^ phosphosite. **(L)** Alphafold3 predicted structure of NUAK1 WT (Uniprot # O60285) with PSD3 isoform 1 WT (Uniprot # Q9NYI0), following the same color scheme. Amino acid side chains are shown for positions −5 to +4 surrounding the PSD3 Ser^476/1011^ phosphosite.

To assess whether affinity purified human PSD3 was phosphorylated by NUAK1, we immunoprecipitated V5-tagged PSD3-WT or phosphomutant V5-PSD3-S476A from HEK293T cells and incubated them with purified NUAK1-WT or NUAK1-KD in an in vitro kinase reaction. We observed robust phosphorylation of WT PSD3 by WT NUAK1, which was completely lost in the PSD3 S476A mutant (Fig.4C-D). We next determined whether NUAK1 and PSD3 interacted with each other. In an affinity pulldown assay with HA-beads, PSD3 was co-immunoprecipitated by HA-tagged NUAK1 but not in the control condition (Fig.4E-F). Similarly, HA-NUAK1 was co-immunoprecipitated by V5-PSD3 (Fig.4G-H), demonstrating that PSD3 and NUAK1 are interacting proteins. To ensure that a co-immunoprecipitating kinase was not responsible for phosphorylating PSD3, we confirmed that bacterially expressed and purified NUAK1 and PSD3 would be sufficient to reconstitute the phosphorylation. We expressed and purified the GST-tagged C-terminal tail of PSD3 which is identical in isoforms 1 and 3 (amino acids 378-513, isoform 3) encompassing the coiled coil domain and C-terminal tail, henceforth referred to as ‘PSD3-CC-Tail’ (Fig.S3B). Incubation of purified NUAK1 WT with PSD3-CC Tail in a kinase activity buffer led to phosphorylation of PSD3-CC-Tail which was abolished in the PSD3 S476A mutant. No phosphorylation was observed as expected with the kinase dead NUAK1 (Fig.4I and 4J).

Next, we used Alphafold3 structure prediction to understand the mechanism through which NUAK1 binds its substrates. First, we performed a prediction of NUAK1 in complex with its known substrate MYPT1, along with the requisite adaptor Protein Phosphatase 1 beta (PP1β) (*48*). In this structure, the region of MYPT1 from amino acids 435-455 (shown as sticks), centered around the phosphosite at Ser^445^, docks into the kinase domain of NUAK1, which is shown as an electrostatic surface model (Fig.4K). The phospho-acceptor Ser^445^ is located adjacent to the binding site for ATP, while basic residues at the −3 and −4 position bind to acidic patches in the NUAK1 kinase domain. The aliphatic isoleucine residues at −5 and +4 positions in MYPT1 dock into small pockets within the NUAK1 kinase domain. To determine whether NUAK1 had a similar binding mode towards the PSD3 phosphosite, we next modeled NUAK1 in complex with full length PSD3 isoform 1 (Fig.4L). Here, the region surrounding the PSD3 Ser^476^ isoform3 (or Ser^1011^ isoform1) phosphosite docks into the catalytic cleft of NUAK1, in a manner highly similar to MYPT1. The phospho-acceptor Ser^476^ of PSD3 is docked directly next to the ATP-binding site, and the basic residues at positions −4 and −3 are aligned with the acidic patches of the NUAK1 kinase domain. Finally, the aliphatic leucine residues at positions −5 and +4 fit into the pockets in the NUAK1 kinase domain in a similar manner to the respective residues in MYPT1. Altogether, our computational modeling highlights the similarity of the binding conformation of NUAK1 towards its known substrate MYPT1 and with PSD3 discovered in our chemical genetic screen.

### PSD3 Phosphorylation at Ser^476^ Regulates ARF6 Activation

Localization of PSD3 to the plasma membrane is required for conversion of GDP-bound ARF6 to active GTP-bound ARF6 (*45*). The C-terminal tail of PSD3 containing both the PH domain and the coiled-coils is required for its plasma membrane association (*45*). Because the NUAK1 phosphosite on PSD3 Ser^476^ lies in the C-terminal tail, we tested whether PSD3 phosphorylation regulates ARF6 localization or activation. Expression of GFP-tagged wildtype PSD3 along with mScarlet-tagged wildtype ARF6 led to the recruitment of ARF6 to the membrane (Fig.5A, top row). We utilized a catalytically dead PSD3 mutant E114K, which has previously been shown to abolish the ability of PSD3 to catalyze the exchange of GDP bound to ARF6 (*49*). Residue Glu^114^ lies within the glutamic finger of the catalytic Sec7 domain, which is responsible for destabilizing GDP and Mg^2+^ to promote nucleotide exchange, and mutation of this glutamic acid to lysine abolishes GEF activity. As expected, PSD3 mutant E114K was unable to recruit and activate ARF6 (Fig.5A, second row). Phosphomutant PSD3 strongly recruited ARF6 to the plasma membrane and its expression led to an aberrant accumulation of large intracellular vesicles positive for both ARF6 and PSD3 (Fig.5A, third row). Because constitutively active ARF6 mutant Q67L leads to similar accumulation of GTP-bound ARF6 that is unable to recycle back^51^, we hypothesized that S476A phosphomutant PSD3 causes aberrant ARF6 activation or prevented the conversion of ARF6 GTP back to its GDP form. To test this hypothesis, we tested whether dominant negative (DN) ARF6 mutant T44N would rescue the effect of PSD3 S476A mutant. Indeed, co-expression of PSD3 S476A along with DN ARF6 did not cause internal accumulation of PSD3 and ARF6 positive vesicles (Fig.5A, bottom row). The area of internalized PSD3/ARF6 positive vesicles increased from 9.6% in WT-PSD3/WT-ARF6 expressing cells to 20.9% in S476A-PSD3/WT-ARF6 expressing cells (n=26 from three biological replicates, p<0.005, one-way ANOVA with Tukey’s multiple comparisons) (Fig.5B). Expression of PSD3 S476A with DN-ARF6 lead to almost complete rescue to 2.2% vesicle area (n=20 from three biological replicates, p<0.0001, one-way ANOVA with Tukey’s multiple comparisons) similar to the inactive E114K PSD3 expression.

**Figure 5.**
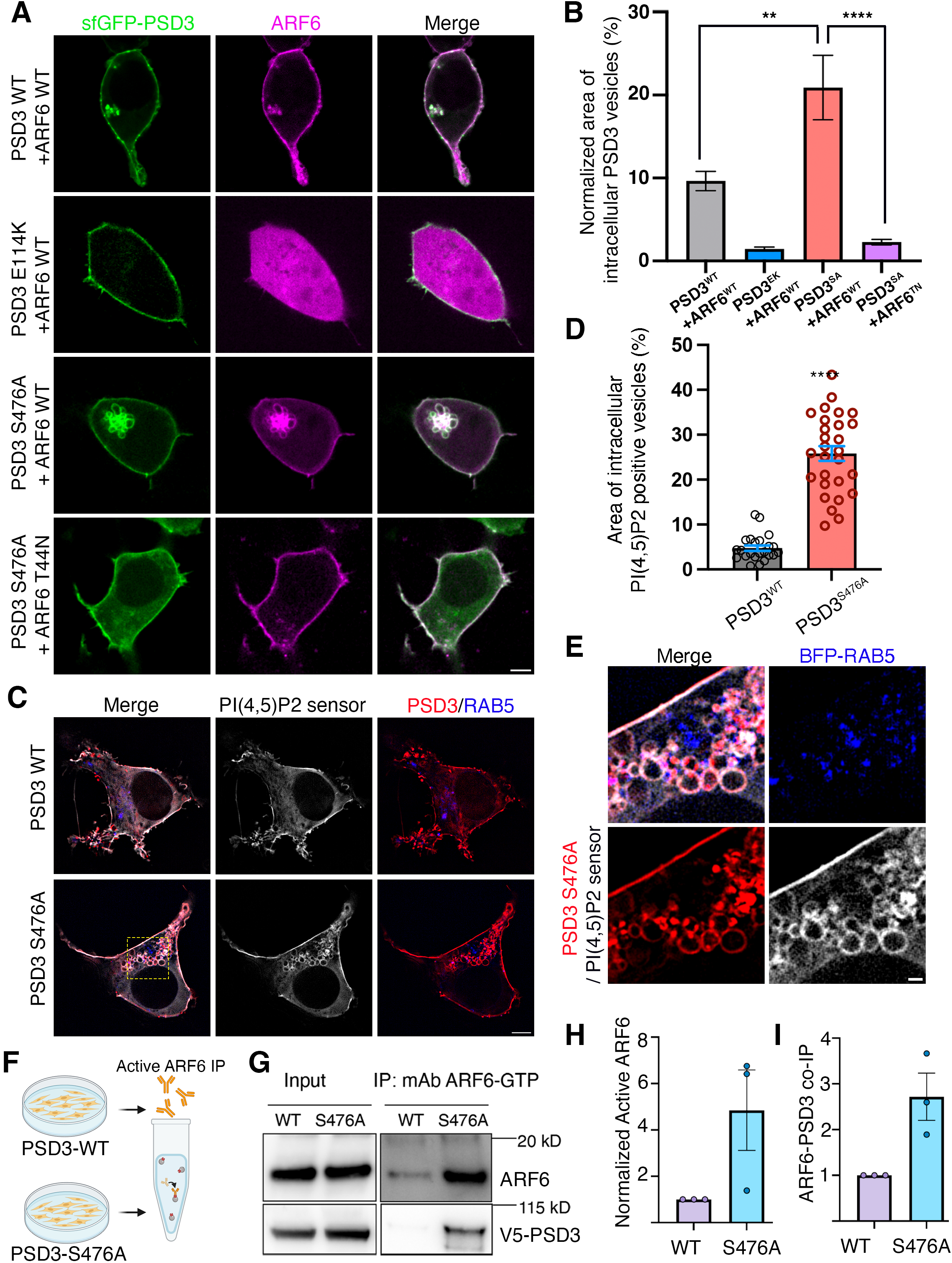
Phosphorylation of PSD3 at Ser476 regulates ARF6 dependent trafficking. **(A)** HEK393T cells co-expressing mScarlet-tagged WT or dominant negative T44N ARF6 (magenta) and sfGFP-tagged WT-PSD3, Sec7 mutant E114K or phosphomutant S476A (green). Scale bar is 5µm. **(B)** Area of intracellular vesicles positive for ARF6 and PSD3 normalized to the total cell area is plotted for each condition (n>20 cells from 3 biological replicates, ** denotes p<0.005 and **** indicates p<0.0001, one-way ANOVA with Tukey’s multiple comparisons). **(C)** HEK393T cells co-expressing the PI(4,5)P_2_ sensor GFP-PLC-delta-PH (gray) along with either WT or phosphomutant PSD3 S476A (red). Scale bar is 5µm. **(D)** Area of intracellular vesicles positive for PI(4,5)P_2_ sensor PH-PLCD1 normalized to the total cell area is plotted for each condition (n>23 cells from 3 biological replicates, **** indicates p<0.0001, unpaired t-test with Welch’s correction). **(E)** Magnified view of inset in panel 5C highlighting intracellular vesicles observed in cells co-expressing the PI(4,5)P_2_ sensor GFP-C1-PLC-delta-PH (gray) along with phosphomutant PSD3 S476A (red) and Rab5-BFP (blue). Scale bar is 1µm. **(F)** Schematic showing the active ARF6 pulldown assay. **(G)** Western blot of active GTP-bound ARF6 immunoprecipitated from HEK293T cells expressing V5-tagged PSD3 WT or S476A. Top blot shows total ARF6 levels in input and IP, while bottom blot shows V5 levels in input and IP. **(H)** Quantification of ARF6 immunoprecipitation from ARF6-GTP immunoprecipitation (n=3 independent experiments, * denotes p<0.05, One-tailed Mann-Whitney test). **(I)** Quantification of V5 immunoprecipitation from ARF6-GTP immunoprecipitation in F (n=3 independent experiments, * denotes p<0.05, One-tailed Mann-Whitney test). All error bars are standard error of the mean.

GTP-bound ARF6 activates phosphatidylinositol 4-phosphate 5-kinase (PIP 5-kinase) that generates phosphatidylinositol 4,5-bisphosphate (PI(4,5)P_2_), which is required for ARF6-dependent endocytosis (*50*). Expression of a GTP hydrolysis-resistant ARF6 mutant or overexpression of PIP 5-kinase induces the formation of PI(4,5)P2 containing actin-coated vacuoles that are unable to recycle membrane back to the plasma membrane (*50*). These vesicles have been shown to contain ARF6-dependent cargo that remains trapped as they are unable to recruit early endosome proteins RAB5 (Ras-related protein Rab5) and EEA1 (early endosome antigen 1) to recycle back to the surface (*51*). To further test our hypothesis that loss of C-terminal PSD3 phosphorylation leads to aberrant ARF6 activation, we determined if levels of PI(4,5)P_2_ and recruitment of RAB5 to internalized membrane would be altered in cells expressing S476A mutant. We expressed WT or S476A phosphomutant PSD3 in HEK293T cells along with the PI(4,5)P_2_ sensor GFP-C1-PLCdelta-PH (*50*) and BFP-tagged RAB5. In cells expressing WT-PSD3, PI(4,5)P_2_ was normally localized to the cell membrane. However, cells expressing phosphomutant PSD3 developed numerous large internal vacuoles that were positive for PSD3 and PI(4,5)P_2_ but devoid of early endosome marker RAB5 (Fig.5C). The total area of internalized vesicles positive for PI(4,5)P_2_ increased six-fold from 4.7% in WT-PSD3 expressing cells to 25.8% in cells expressing the S476A mutant PSD3 (n=23 cells from three biological replicates, p<0.0001, unpaired t-test with Welch’s correction) (Fig.5D). Visualization of single z frame images of internalized PIP_2_ vesicles through super resolution microscopy(*52*) indicated clear absence of RAB5 on the PSD3 / PI(4,5)P_2_ positive vesicles (Fig.5E).

To biochemically assess whether ARF6 activation is increased in cells expressing PSD3 S476A, we expressed V5-tagged PSD3 WT or S476A in HEK293T cells and immunoprecipitated active ARF6 from these cells using an antibody against the GTP-bound form of active ARF6. Active ARF6 was affinity isolated from transfected cells using the active ARF6 antibody (Fig.5F) and total ARF6 levels were determined by with another ARF6 antibody that recognizes ARF6 regardless of nucleotide state. We observed a 4.5-fold increase in active ARF6 pulldown from cells expressing PSD3 S476A compared with PSD3 WT (Fig.5G-H). Further, we observed a significant increase in the interaction between ARF6 and PSD3 in the phosphomutant compared to WT-PSD3 (Fig.5I). This suggests a negative regulatory mechanism where phosphorylation of PSD3 at Ser^476^ by NUAK1 decreases PSD3 association with ARF6 thereby limiting further ARF6 activation. Together, these results demonstrate that NUAK1 mediated phosphorylation of PSD3 at Ser^476^ is important for regulating ARF6 activation and ARF6-mediated endosomal trafficking.

### NUAK1 Phosphorylation of PSD3 Regulates Dendritic Spine Maturation

Next, we investigated the role of PSD3 phosphorylation in neuronal development. *PSD3* is widely expressed in the central nervous system both during and after developmental time periods. In mouse brain, the highest expression is within the hippocampus and cerebral cortex (Fig.S4A), while in the human brain *PSD3* expression is broad including hippocampus, cortex, striatum, and cerebellum. *PSD3* expression increases during embryonic brain development, peaks at birth and is maintained at high levels throughout the human lifespan (Fig.S4B). PSD3 through its function as a GEF for ARF6 (*44*), has important roles in dendritic spine development (*53*, *54*). For instance, expression of constitutively active, fast cycling mutant of ARF6, T157A, significantly increases synaptic maturation such that density of mature spines positive for PSD95 increases while filopodial protrusions are decreased (*54*). Further, ARF6-mediated endocytic recycling also can directly influence dendritic spine maturation through endocytosis mediated internalization of adhesion molecules such as telencephalon (*55*). To determine how NUAK1-mediated phosphorylation of PSD3 impacts neuronal development, we transfected rat hippocampal neurons with membrane marker myr-tdTomato with either GFP-tagged wildtype PSD3 or the S476A phosphomutant and imaged them after 48hr at DIV14. WT PSD3 localized throughout the neuron including soma, dendrites, and axons, but was particularly enriched in mushroom-shaped spines (Fig.6A-B). Notably, we observed that neurons expressing the phosphomutant PSD3 S476A showed significantly increased enrichment of PSD3 within dendritic spines concomitant with increased number of mushroom spines (Fig.6C, Fig.S4C). The percentage of dendritic spines enriched in PSD3 increased from a mean of 33.8% (n=16 from three biological replicates) in WT PSD3 expressing neurons to 63.8% (n=17 from three biological replicates, p<0.0001, t-test with Welch’s correction) in neurons expressing the S476A phosphomutant (Fig.6C). To further analyze the impact of PSD3 phosphorylation on dendritic protrusions, we binned protrusions into four classes: stubby, thin, filopodia, and mushroom. While there were no significant differences in the percent of stubby and thin protrusions, the filopodial protrusions decreased from 13.9% to 7.5% in neurons expressing the phosphomutant PSD3 S476A (n=56 neurons from three biological replicates, SEM=1.33, p<0.05, t-test with Welch’s correction) and mushroom spines increased from 36.1% in WT (n=65 from three biological replicates, SEM=2.08) to 48% in S476A mutant expressing neurons (n=56 from three biological replicates, SEM=2.0, p<0.0001, t-test with Welch’s correction) (Fig.S4C). No significant differences were observed in density of protrusions between the wildtype and S476A mutant PSD3 expressing neurons (Fig.S4D). These results suggest that NUAK1 mediated phosphorylation of PSD3 at Ser^476^ does not impact formation of new protrusions but rather controls the maturation of filopodial protrusions into mature mushroom shaped spines enriched in PSD3. To assess whether morphological maturity of spines led to functional changes, we measured the density of postsynaptic protein PSD95. DIV12 hippocampal neurons were transected with PSD3-GFP WT or S476A along with the postsynaptic marker PSD95-RFP and then imaging at DIV14. We observed a significant increase in density of postsynaptic marker PSD95 in neurons expressing S476A phosphomutant compared to neurons expressing WT PSD3 (p<0.0001, unpaired t-test with Welch’s correction).

**Figure 6.**
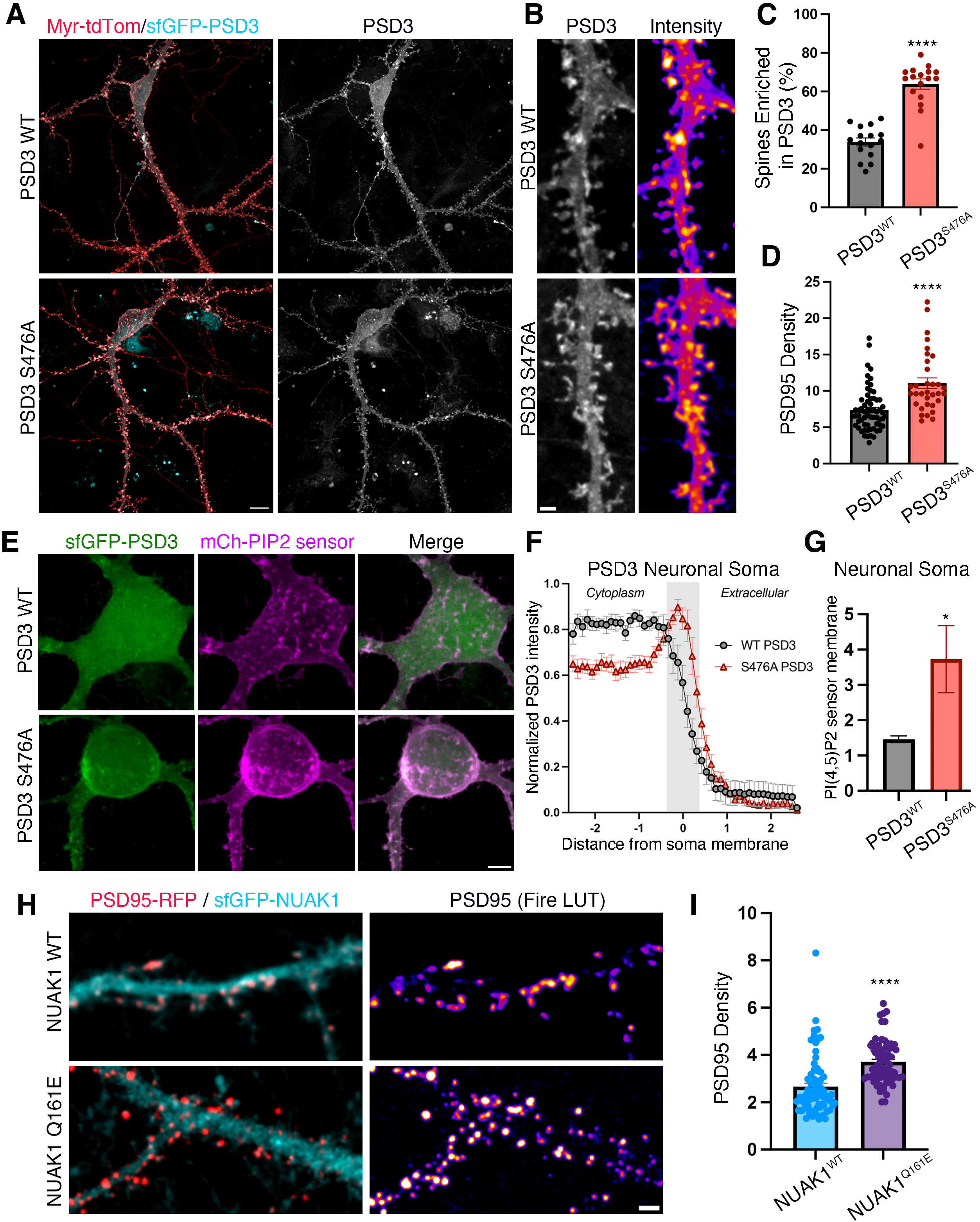
NUAK1 mediated PSD3 phosphorylation at Ser476 restricts maturation of filopodia into dendritic spines. **(A)** Representative images of DIV14 rat embryonic hippocampal neurons co-expressing membrane marker myristoylated-tdTomato (red) along with either sfGFP-tagged wildtype (WT) PSD3 or phosphomutant S476A (cyan). Scale bar is 10μm. **(B)** Magnified insets are shown to highlight the morphology of dendritic spines and the enrichment for PSD3 within spines. Grayscale and pseudo-colored images show intensity of PSD3 distribution in WT and S476A conditions. Scale bar is 1µm. **(C)** Quantification of percent spines enriched in PSD3 in hippocampal neurons expressing WT-PSD3 and S476A phosphomutant (n>15 neurons for each condition from three biological replicates, **** denotes p<0.0001, t-test with Welch’s correction). All error bars are standard error of the mean. **(D)** Quantification of PSD95 density per 10μm dendrite length in hippocampal neurons expressing RFP tagged PSD3 along with GFP tagged WT-PSD3 and S476A-PSD3 (n>30 dendrite segments from at least 10 neurons from three biological replicates for each condition, **** denotes p<0.0001, t-test with Welch’s correction). **(E)** Representative images of DIV14 rat embryonic hippocampal neurons co-expressing PI(4,5)P_2_ sensor mCherry-C1-PLC-delta-PH (magenta) along with either sfGFP-tagged wildtype (WT) PSD3 or phosphomutant S476A (green). Scale bar is 5μm. **(F)** Distribution of normalized sfGFP-PSD3 intensity as a function of distance from the plasma membrane of the neuronal soma is plotted (n=10 neurons from three independent experiments). **(G)** Normalized intensity of PI(4,5)P_2_ sensor intensity at the soma membrane in neurons expressing WT or S476A PSD3 is plotted (n=10 neurons from three independent experiments, p<0.05, t-test with Welch’s correction). **(H)** Representative images of DIV14 rat embryonic hippocampal neurons co-expressing postsynaptic marker PSD95-RFP (red) along with either sfGFP-tagged WT-NUAK1 or ASD variant Q161E-NUAK1 (cyan). Scale bar is 2μm. **(I)** Quantification of PSD95 density per 10μm dendrite length in hippocampal neurons expressing RFP tagged PSD3 along with GFP tagged WT-NUAK1 and Q161E mutant NUAK1 (n>65 dendrite segments from at least 20 neurons from three independent experiments for each condition, **** denotes p<0.0001, t-test with Welch’s correction). All error bars are standard error of the mean.

To determine whether PSD3 phosphorylation in neurons impacted ARF6 activation in neurons, we measured levels of PI(4,5)P2 levels in neurons expressing either WT or S476A mutant PSD3 using the sensor Scarlet-C1-PLCdelta-PH (Fig.6E). Neurons expressing the GFP-PSD3 S476A mutant showed increased accumulation of the PSD3 at the membrane concomitant with a significant increase in the intensity of the PI(4,5)P2 at the neuronal soma membrane (Fig.6F-G) (n=10 neurons, p<0.05, unpaired t-test with Welch’s correction). Next, we tested whether NUAK1 kinase-deficient ASD mutant Q161E would impact spine maturation similar to phosphomutant PSD3. Rat embryonic hippocampal neurons were transfected with synaptic marker PSD95-RFP along with GFP-tagged NUAK1 WT or Q161 mutant on DIV12 and density of PSD95 punctae was measured at DIV14. Similar to the effect of expressing phosphomutant PSD3, we observed a significant increase in the density of PSD95 in neurons expressing the catalytically deficient NUAK1 compared to WT-NUAK1 (Fig.6H-I) (p<0.0001, n>65 from three experiments, unpaired t-test with Welch’s correction). Together, our results suggest that NUAK1-mediated phosphorylation of PSD3 at Ser^476^ controls ARF6 activation and dendritic spine maturation in neurons.

## Discussion

We utilized a chemical-genetic screen to reveal phosphorylation targets of the autism-associated kinase NUAK1 in the brain. This unbiased approach allowed us to map the basophilic consensus motif preferred by NUAK1. We uncovered direct substrates of NUAK1 in cellular processes NUAK1 has been previously implicated in including RNA splicing, axon growth, and dendrite development, as well as discovered new functions of NUAK1 in regulation of endocytosis and dendritic spine and synapse maturation (Fig.7A). Among the identified substrates, we focused on the ARF6-specific guanine nucleotide exchange factor PSD3 and mechanistically delineated how its phosphorylation by NUAK1 negatively controls ARF6-dependent endocytosis and dendritic spine maturation (Fig.7B and 7C).

**Figure 7.**
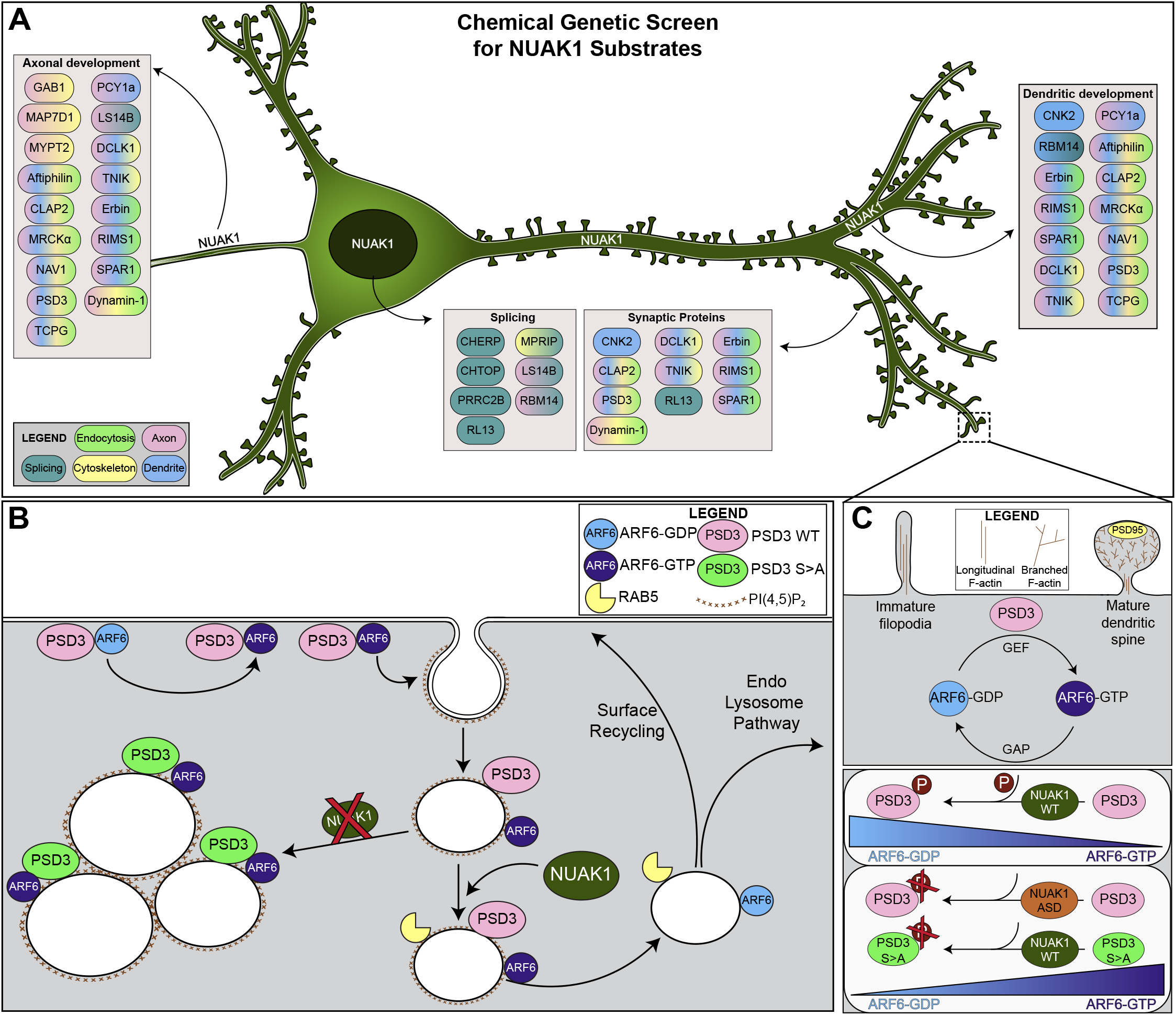
NUAK1 phosphorylation targets in neuronal development. **(A)** Schematic shows substrates of NUAK1 identified by the chemical genetic screen which are binned by their roles in axonal, dendritic, and synaptic development, as well as splicing. Each substrate is color-coded according to its known function(s), with light green for endocytosis, teal for splicing, yellow for cytoskeleton, pink for axon, and blue for dendrites, with the legend in the lower left panel showing each individual color. Substrates with multiple functions are indicated with gradients of 2-4 colors. **(B)** Schematic representation of the role of PSD3 and NUAK1 in ARF6 activation. At the plasma membrane, PSD3 recruits inactive, GDP-bound ARF6. PSD3 exchanges GDP for GTP, generating active ARF6, which promotes the formation of PI(4,5)P_2_. This increase in PIP_2_ causes internalization of PSD3 and ARF6 on PIP_2_-enriched vesicles. In normal conditions, Rab5, ARF6 GAPs, and lipid phosphatases are recruited to these vesicles, leading to a cessation of ARF6 activity and depletion of PIP_2_. These vesicles can then continue in the endolysosomal pathway or be recycled to the cell surface. When NUAK1 is not present, or when the PSD3 Ser^476^ phosphosite is mutated to alanine, ARF6 remains active and bound to GTP, accumulating in large, PIP_2_-enriched vesicles. **(C)** Schematic of the role of PSD3 in dendritic spine maturation. PSD3 activation of ARF6 leads to an increased proportion of mature, mushroom-shaped dendritic spines and a corresponding loss of immature filopodia. In normal conditions, PSD3 is phosphorylated by NUAK1, keeping ARF6 activation in check. When NUAK1 is inactive or when the PSD3 phosphosite is mutated to alanine, PSD3 phosphorylation is lost, leading to an aberrant increase in the proportion of active ARF6.

We report the biochemical characterization of autism-associated NUAK1 variants and show that NUAK1 mutations have differential impact on catalytic activity and subcellular localization. The residue Gln^161^ in the αE-helix in the C-lobe is under high evolutionary constraint and makes key contacts with residues Asn^191^ and Ile^192^ in the β8 strand that stabilize the catalytic loop of NUAK1. We found that the autism variant Q161E led to a significant decrease in the catalytic activity of NUAK1. The corresponding Gln residue in protein kinase A (PKA) is Gln^149^ which has been shown to allosterically regulate the DFG loop during transitions between catalytic active and inactive states (*56*, *57*). The premature termination variant Q433* is predicted to escape nonsense mediated decay as this residue is encoded in the final exon of NUAK1. Further, our data show that Q433* has intact catalytic activity. Upon expression in neurons and dividing cells, however, we found that the truncated NUAK1 Q433* is trapped in nuclear speckles positive for SC35. The third NUAK1 disease variant, A653V, also does not impact catalytic activity but decreases NUAK1 association with nuclear speckles. Our structural analyses suggest that the residue Ala^653^ lies in an amphipathic helix within the C-terminal tail. Mechanisms through which the latter two variants have opposing impacts on NUAK1 association with nuclear speckles remains to be determined. Collectively, our data suggest that heterozygous ASD-associated mutations in NUAK1 are predicted to either decrease substrate phosphorylation or alter substrate availability by shifting the localization of the active kinase between cytoplasm and nuclear speckles, two distinct locations where NUAK1 likely phosphorylates non-overlapping set of substrates. Indeed, our dataset of identified NUAK1 substrates includes both nuclear and cytoplasm enriched proteins. It is important to note that *NUAK1* is widely expressed in human brain and temporally much earlier during brain development compared to mice. Transcriptomic analyses show emergence of NUAK1 expression in neuronal progenitors which continues post differentiation while in mice *Nuak1* expression is mostly limited to post mitotic neurons (*58*). Therefore, NUAK1 nuclear phosphorylation substrates may have a more prominent role in human brain development than in mice.

To shortlist the putative substrate candidates identified in our chemical-genetic screen, we used the following criteria, reasoning it would be true for substrates directly phosphorylated by NUAK1: (1) the substrate is phosphorylated by NUAK1 in vitro, (2) mutation of the identified phosphosite abolishes substrate phosphorylation, (3) substrate biochemically interacts with NUAK1, and that (4) purified NUAK1 and substrate in vitro would be sufficient to reconstitute phosphorylation at the identified phosphosite. PSD3, an ARF6-specific GEF identified in our screen, meets all four criteria consistent with a bona fide NUAK1 substrate. Human PSD3 isoform 3 is phosphorylated at Ser^476^ (corresponding to Ser^1011^ isoform 1), an evolutionarily conserved site that fits the NUAK1 consensus motif. Notably, mutation of the Ser^476^ residue in PSD3 to alanine abolishes all phosphorylation by NUAK1, indicating it is the only site within PSD3 that is phosphorylated by NUAK1. The phosphosite is situated in the C-terminal tail of PSD3 following the PH domain which binds PI(4,5)P_2_ and the coiled coil region which binds cortical actin. The C-terminus of PSD3 mediates its plasma membrane association which is required for activation of ARF6 at the membrane via the catalytic Sec7 domain. While it is unlikely that phosphorylation at Ser^476^ directly impacts Sec7 domain-mediated GEF activity, our data shows that phosphomutant PSD3 S476A is more strongly associated with GTP-bound ARF6. Accumulation of PI(4,5)P_2_ positive vesicles in cells expressing phosphomutant PSD3 are phenotypically similar to those observed upon expression of the ARF6 Q67L mutant which cannot hydrolyze GTP. Another possibility is that phosphorylation of the PSD3 tail does not impact ARF6 activity but prevents recruitment of ARF6 GAPs that convert GTP-bound ARF6 to GDP. Our immunoprecipitation results demonstrate that PSD3 S476A significantly increases the proportion of active GTP-bound ARF6 in HEK293T cells, and that this mutant shows increased association with this active form of ARF6. This suggests that phosphorylation of PSD3 at Ser^476^ by NUAK1 serves as a regulatory mechanism to decreases PSD3 mediated activation of ARF6.

It is important to note that copy number variation, missense variants, and frameshift mutations in PSD3 have been identified in patients with autism spectrum disorder (*29*, *30*, *59*). ARF6 GTPase is an established regulator of actin dynamics and membrane trafficking, and plays an essential role in neurite development, axon growth, dendritic spine maturation, and synaptic plasticity (*60*). PSD3 is enriched within dendritic spines, and our data shows that abolishing NUAK1 mediated phosphorylation at Ser^476^ further increases PSD3 spine enrichment. This also causes a loss of filopodial protrusions and stabilization of mature mushroom dendritic spines and leads to an increase in synaptic density. This is consistent with the impact of expressing the fast-cycling ARF6 mutant T157A, which increases the conversion of filopodia into mature spines (*54*). The impact of PSD3 variants associated with autism spectrum disorder on spine stability, maturation, and ARF6 activity has not been tested and will be subject of future investigation. In addition to autism, both NUAK1 and PSD3/ARF6 have been implicated in cancer progression, fatty liver, and metabolic disease (*19*, *61–65*). Our study raises the intriguing possibility that NUAK1-mediated phosphorylation of PSD3 underlies diverse physiological effects of ARF6-dependent endocytic trafficking.

## Methods

### Expression Constructs, Cloning and Reagents

Full-length human NUAK1 was PCR amplified from K84M kinase-DEAD Nuak1-3x Flag (Addgene #97219) and inserted into vector sfGFP-C1 (Addgene #54579) or PRK5-HA (PMID: 28065648) using PCR amplification followed by Gibson Assembly (NEB). The K84M mutation was corrected using primers to generate WT NUAK1. All mutants of NUAK1 (K84M, M132A, M132G, Q161E, Q433*, A653V) were generated using PCR amplification followed by Gibson assembly (NEB). NUAK1 Q433* was cloned into the bacterial expression vector 8xHis-MsyB (HM) (Addgene #112570) to generate HM-NUAK1 1-433 using PCR amplification followed by Gibson assembly (NEB). The K84M or M132G mutations were introduced in this plasmid using PCR amplification followed by Gibson assembly (NEB). Full-length human PSD3 isoform 1 (canonical) was obtained from DNASU and inserted into an N-terminal V5 mammalian expression using Gateway cloning (Thermo). PSD3 isoform 3, which is 513 amino acids long, shares all except the first 11 amino acids with isoform 1. PSD3 isoform 3 was cloned using PCR amplification to insert the 11 amino acids at the N-terminus, followed by Gibson assembly (NEB). PSD3-isoform 3 E114K and S476A point mutations were cloning using PCR amplification followed by Gibson assembly (NEB). V5-PSD3-iso3-sfGFP was cloned by inserting sfGFP from sfGFP-C1 vector (Addgene #54579) at the C-terminus of PSD3 followed by a stop codon using PCR amplification and Gibson Assembly. mCherry-V5-PSD3-iso3 was cloned by inserting the mCherry from mCherry-Rab5 (Addgene #49201) upstream of V5-PSD3-iso3 using PCR amplification followed by Gibson Assembly (NEB). GST-PSD3-coiled coil tail was generated by PCR amplification of amino acids 378-513 of PSD3-isoform 3 followed by Gibson Assembly (NEB) with PCR-amplified pGex-4T3 vector (Addgene #31669). Arf6-mNeonGreen was obtained from Addgene, and the mNeonGreen was exchanged for mScarlet from mScarlet-Rab7 (Addgene #169068) using PCR amplification and Gibson Assembly (NEB). Arf6-mScarlet T44N (DN) mutation was introduced using PCR amplification followed by Gibson Assembly (NEB).

### Cell Line Maintenance and Transfection

HEK293T (human embryonic kidney 293 SV40 large T antigen) cells (RRID: CVCL_0063) were grown in Dulbecco’s modified Eagle’s medium (Thermo Fisher Scientific, Gibco) with 10% fetal bovine serum (Axenia BioLogix) and 1% penicillin-streptomycin (Invitrogen). Cells were maintained at 5% CO_2_ and 37°C and passaged every 3 to 4 days. Cells were tested for mycoplasma using the ABM PCR-based kit. For biochemical experiments, HEK293T cells were transfected with 0.5-1 ug plasmid DNA using the JetOptimus system (Genesee Scientific) following the manufacturer’s instructions. For confocal imaging, HEK293T cells were transfected with 100-500 ng plasmid DNA using Lipofectamine 2000 and OptiMEM (Thermo Scientific) according to the manufacturer’s instructions.

Primary neurons were obtained from dissociating hippocampi obtained from embryonic day 18 (E18) Sprague-Dawley rat (Envigo; RRID: MGI:5651135). Animal work was performed under University of Washington Institutional Animal Care and Use Committee approved protocol 4421-01. Rat embryos were trypsin-dissociated and plated at a density of 150,000 neurons per 18-mm glass coverslip (Thermo Fisher Scientific) or 450,000 neurons per 35-mm glass-bottom dish (MaTek), which were coated with poly-D-lysine (0.06 mg/ml; Sigma-Aldrich) and laminin (0.0025 mg/ml; Sigma-Aldrich). Plating medium containing heat-inactivated 10% fetal bovine serum (HyClone), 0.45% dextrose, sodium pyruvate (0.11 mg/ml), and 2 mM glutamine in minimum essential medium with Earle’s Balanced Salt Solution was used. Medium was changed 4 hours after seeding neurons into maintenance medium consisting of Neurobasal medium (Invitrogen), 2% B27 (Invitrogen), penicillin (50 U/ml) and streptomycin (50 μg/ml), and 0.5 mM Glutamax (Thermo). Half of the medium was replaced with new maintenance medium every 3 to 4 days. For exogenous gene expression, neurons were transfected with 0.5 to 1.0 μg of plasmid DNA per coverslip in a 12-well plate (or 1 to 2 μg of DNA per 35-mm dish) using Lipofectamine 2000 (Invitrogen) following the manufacturer’s guide.

WTC11 induced pluripotent stem cells (iPSCs; Corielle Institute; RRID: CVCL_Y803) were differentiated into neural progenitor cells (NPCs) as reported previously (*66*). Stem cells were treated with 1U/mL Dispase, lifted using a cell scraper, resuspended in mTeSR media (STEMCELL Technologies) containing 10μm ROCK Inhibitor (Selleck Chemical), and then transferred to uncoated T25 flasks for embryoid body (EB) formation. After 24–48 hours (Day 0), EBs were gravity settled and transferred to new uncoated T25 flasks in neural media (DMEM/F12 (Gibco), 1% N-2 supplement (Gibco), 1% Non-essential amino acids (Gibco), 2μg/mL Heparin, 1% Penicillin/Streptomycin [Gibco]) containing small molecule TGF-β/SMAD inhibitors, SB431542 (5μM; Peprotech) and LDN193189 (0.25μM; Peprotech). On Day 3, EBs were transferred to Matrigel (1:50 dilution; Corning)-coated 6 well plates in fresh neural media with SB and LDN. Media was changed with neural media without SB and LDN every other day or as needed until Day 11 for formation of neural rosettes. On Day 11, neuroepithelia were mechanically lifted using a cell scraper and transferred to uncoated T25 flasks in neural media. Neurospheres were fed with neural media every other day, or as needed, until Day 25. On Day 25, neurospheres were plated onto Matrigel-coated plates in STEMDiff™ Neural Progenitor Media (STEMCELL Technologies) and NPCs were obtained. NPCs were tested for neural progenitor markers Nestin (Invitrogen) and PAX6 (Sigma).

### Protein Purification

#### HM-NUAK1 purification

NUAK1 (WT, KD, or M132G) were inserted into 8x-His-tagged pExp-MsyB (His-MsyB, or HM) vector for bacterial expression and purification. Due to poor expression of full-length NUAK1 1-661, we elected to use NUAK1 1-433, which is kinase active but contains less unstructured domains than full-length NUAK1. Expression of NUAK1 1-433 was much improved over full length, so we moved forward with purification of HM-NUAK1 1-433 WT, K84M, and M132G. HM-NUAK1 1-433 WT, K84M, or M132G were transformed into BL21 *E. coli* (New England Biolabs) and sequenced using full-plasmid sequencing (Plasmidsaurus) to confirm the sequences were correct. 50 mL cultures of HM-NUAK1 were grown overnight with Carb and used to inoculate 1L cultures, which were grown at 37C with 220 rpm shaking until Optical Density (OD) of 0.6-0.8 was reached. At this point, the incubator was cooled to 20C (room temperature), and IPTG (Isopropyl β-d1-thiogalactopyranoside) was added to all cultures for a final concentration of 0.4mM. Protein induction was performed overnight at 20C with 220 rpm shaking. Bacteria were pelleted the next day by spinning at 3,200 rcf at 4C for 15 minutes, followed by removal of LB and addition of 30 mL 1x DPBS. The cultures suspended in PBS (Phosphate buffered saline) were transferred to 50 mL conical tubes and spun at 3,200g at 4C for 15 minutes. The PBS was then removed, and 25 mL lysis buffer (20 mM Tris pH 7.5, 200 mM NaCl, 0.4 mM TCEP, 20 mM imidazole, 1x cOmplete protease inhibitor (Sigma), 0.1% PMSF, and 10% glycerol) was added to the tube. Bacteria were lysed using sonication at 25% power for 3.5 minutes, followed by centrifugation at 24,500g for 60 minutes at 4C to pellet debris. Following centrifugation, the supernatant was added to 1 mL HisPur Ni-NTA resin (Thermo) for 60 min at 4C with rotation to bind HM-NUAK1 to the beads. After binding, beads were added to a fritted column and the supernatant allowed to drain by gravity. The beads were then washed with 20 mL lysis buffer, followed by the addition of lysis buffer + 200 mM imidazole. This high imidazole buffer was incubated with the beads for 5 minutes, after which eluted protein was collected every 10 minutes for a total of 6 elutions, and quantified using Pierce 600nm protein assay (Thermo). The eluted protein was then incubated with sulphopropyl (SP) beads (GE healthcare) to enrich for acidic proteins for 1 hour rotating at 4C, followed by elution with 450mM NaCl. Finally, proteins were concentrated to a final concentration of ∼10 ug/mL using concentrators with a 30kD molecular weight cutoff (Thermo). Following concentration, protein aliquots were flash frozen in liquid nitrogen and stored at −80C. All purification samples were assayed for purity by SDS-Page, Coomassie staining, and Western blotting with anti-His-HRP antibody (Proteintech).

#### GST-PSD3 purification

PSD3-CC tail (WT or S>A) was cloned into the pGEX-4T3 bacterial expression vector, transformed into BL21 *E. coli*, and sequenced using full-plasmid sequencing (Plasmidsaurus). 25 mL overnight cultures of GST-PSD3 were grown overnight with Carbenicillin and used to inoculate 1L cultures, which were grown at 37C with 220 rpm shaking until Optical Density (OD) of 0.6-0.8 was reached. At this point, the incubator was cooled to 30C, and IPTG was added to all cultures for a final concentration of 0.3mM. Protein induction was performed for 2 hours at 30C with 220 rpm shaking. Bacteria were pelleted by spinning at 3,200 rcf at 4C for 15 minutes, followed by removal of LB and addition of 30 mL 1x DPBS. The cultures suspended in PBS were transferred to 50 mL conical tubes and spun at 3,200g at 4C for 15 minutes. The PBS was then removed, and samples were resuspended in 7 mL lysis buffer (50mM Tris pH 8.0, 200mM NaCl, 5mM EDTA, 10% glycerol, 5mM DTT, 1x cOmplete protease inhibitor (Sigma), 0.1% PMSF). 4 mg of lysozyme was added to each sample, followed by incubation for 30 minutes on ice with occasional mixing. After this step, Triton was added to the sample for a final concentration of 1%, followed by lysis with sonication as described above. After lysis, samples were centrifuged at 24,500g for 60 minutes at 4C to pellet debris. Columns for batch binding were prepared using 500 uL of glutathione resin (Thermo) and 10 mL binding buffer (50mM Tris pH 8.0, 200mM KCl, 1mM DTT). The buffer was removed prior to the addition of the post-centrifugation bacterial supernatant to the beads. The binding of GST-PSD3 to the beads was performed overnight at 4C with rotation. The next day, columns were washed 2x with 15 mL wash buffer (50mM Tris pH 8.0, 200mM KCl, 0.1% Tween-20 1mM DTT), followed by 2x washes with 1 mL binding buffer to remove detergent. Samples were then eluted in binding buffer + 25 mM glutathione in 500 uL fractions every 10 minutes, for a total of 6 elutions. Protein concentration was quantified using Pierce 600nm protein assay (Thermo), and purification samples were assayed for purity by SDS-Page, Coomassie staining, and Western blotting with anti-GST-HRP antibody (Thermo). Following protein quantification, GST-PSD3 was diluted to 1 mg/mL final concentration with binding buffer, and aliquots were flash frozen in liquid nitrogen and stored at −80C.

#### Immunoprecipitation and Kinase Assay

HEK293T cells were transfected with 0.5-1 ug plasmid DNA using the JetOptimus system according to the manufacturer’s instructions (Genesee Scientific). After overnight transfections, cells were lysed in HKT lysis buffer (25 mM HEPES pH 7.2, 150 mM KCl, 1% Triton X-100, 1mM DTT) with 1x cOmplete protease inhibitor (Sigma). Lysate was then spun for 10 minutes at 15,000g at 4C to pellet out cell debris. For pre-conjugated beads (HA, V5), beads were equilibrated with 2x HKT washes, followed by addition of 1 mL cell lysate for 30 uL equilibrated beads. For non-conjugated beads (Protein-G), beads were equilibrated with 2x HKT washes, followed by pre-clearing of 1 mL cell lysate with 15 uL equilibrated beads for 30-60 min at 4C rotating. 1 ug of mouse primary antibody (anti-GFP, Roche) was conjugated to 30 uL of Protein G beads during this pre-clear step, followed by addition of pre-cleared lysate to the conjugated beads for immunoprecipitation. Proteins were immunoprecipitated for 3-16 hours at 4C with rotation. Following IP, samples were washed 2x with HKT, 1x with high salt HKT (25 mM HEPES pH 7.2, 1 M NaCl, 1% Triton X-100, 1mM DTT) for 10 minutes, 2x with HK buffer (25 mM HEPES pH 7.2, 1 M NaCl, 1mM DTT), and 2x with kinase buffer (20 mM Tris HCl pH 7.5, 10 mM MgCl2, 1 mM DTT, 1x cOmplete protease inhibitor). Following bead washes, kinase labeling reaction was performed for 45 minutes at 30C on a shaking heat block (Thermo) in kinase buffer with 0.5mM ATP (Sigma) or ATP S (Thermo) and 2x Halt Phosphatase inhibitor (Thermo). Following the kinase reaction, samples with normal ATP were quenched by the addition of 4X Lammeli-SDS sample buffer (Thermo) with 200mM DTT. Labeling was quenched in samples with ATP S by the addition of EDTA to a final of concentration of 50mM, and thiophosphates were then alkylated with p-nitrobenzyl mesylate (pNBM, Sigma) at a final concentration of 6.67mM for 45 minutes at room temperature. After alkylation of the ATPgS-containing samples, the reaction was quenched by the addition of 4X Lammeli-SDS sample buffer (Thermo) with 200mM DTT. For co-IP experiments, 0.5μg of V5-PSD3 was co-transfected with 0.5μg of HA-NUAK1 WT or HA-empty vector with JetOptimus (Genesee Scientific) as above. This was followed by lysis, equilibration of beads, and immunoprecipitation with either HA or V5-conjugated beads for 3 hours. Following IP, samples were washed 3x with HKT and 3x with HK, followed by addition of 4X Lammeli-SDS sample buffer (Thermo) with 200mM DTT.

#### Microscopy

Live and fixed cell confocal imaging was performed on a Nikon Ti2 Eclipse-CSU-X1 confocal spinning disk microscope equipped with four laser lines—405, 488, 561, and 670 nm—and the Nikon Elements software. An Andor Zyla4.2 Plus sCMOS camera was used for image acquisition. The microscope was caged within the OkoLab environmental control setup, enabling temperature and CO_2_ control during live imaging. Imaging was performed using Nikon 1.49 100×, Apo 60×, or 40× oil objectives. All image analyses were done using the open access Fiji software. The Zeiss ELYRA7 Lattice SIM2 structural illumination microscope was used for super resolution imaging. Live cell imaging was performed on a Zeiss Axio Observer 7-Lattice SIM inverted microscope for super resolution microscopy equipped with four laser lines – 405 nm (50mW), 488 nm (100mW), 561 nm (100mW) and 642 nm (150mW) – and the ZEN imaging software. Two Hamamatsu ORCA-Fusion BT sCMOS cameras were used for simultaneous image acquisition. The microscope was equipped with a Zeiss Stage Top incubation and gas system for Lattice SIM inverted microscopes, enabling temperature, humidity, and CO2 control during live imaging. Imaging was performed using Plan-Apochromat 63x/1.40 oil DIC objective.

#### Immunofluorescence and Western Blotting

All cells, including neurons, were fixed using warm 4% paraformaldehyde with 4% sucrose for 20 min at room temperature followed by three washes with 1x phosphate-buffered saline (PBS). One-hour incubation with blocking buffer [4% normal donkey serum, 0.2 M glycine (pH 7.4), and 0.1% Triton X-100, in 1x PBS] was followed by overnight incubation with primary antibody at 1:100-1000 dilution in blocking buffer at 4°C. After six 5-min washes in PBS, cells were incubated with secondary antibody at 1:1000 dilution in blocking buffer for overnight at 4°C. Coverslips were washed 6 times in PBS and then mounted onto microscope glass slides with Fluoromont-G (EMS) and sealed with clear nail polish (EMS). Samples for Western blot analyses were treated with NuPAGE LDS sample buffer (4x) (Invitrogen) with Dithiothreitol (DTT) at 200mM and subsequently heated for 10 min at 95°C. Samples were electrophoresed on NuPAGE 4 to 12% Bis-Tris polyacrylamide gels (Invitrogen) with NuPAGE MOPS or MES running buffer (Invitrogen). Western blots were transferred to Immobilon-P (MilliporeSigma) with transfer buffer [25 mM Tris, 192 mM glycine, 0.1% SDS and 20% (v/v) methanol]. Transferred blots were blocked in 5% nonfat dry milk or 5% Bovine Serum Album (Equitech Bio) and subjected to primary antibody overnight, washed 3x with blocking buffer, and incubated 3 hours at room temperature with horseradish peroxidase–conjugated secondary antibody before visualization with Pierce SuperSignal West Pico PLUS Chemiluminescent Substrate (Thermo Fisher Scientific). Western blot images were obtained using the ChemiDoc Imaging System (Bio-Rad). Western blot images were analyzed and quantified using Image Lab Software for Mac Version 6.1 (Bio-Rad).

#### Mouse Brain Lysate

To prepare mouse brain lysate, P13-P15 C57BL/6 (RRID: MGI:2159769) mice pups of both sexes were euthanized by CO2 (5psi for 10min) followed by decapitation. Eight mice were utilized and were not sexed prior to sacrifice. The brains were dissected out and incubated in high detergent lysis buffer (20mM Tris pH7.5, 100mM NaCl, 10mM MgCl2, 0.5mM DTT, 1% TritonX-100, 1% deoxycholic acid, and 1x cOmplete protease inhibitor) on ice for 30min. Homogenization was achieved by sonication. Supernatant was collected after centrifugation at 14,000g at 4°C for 10 min. Supernatants from all tissues were pooled together and aliquots were prepared by diluting with lysis buffer without detergent to 20mg/ml, followed by flash freezing in liquid nitrogen and storage at −80°C.

#### NUAK1 Substrate Labeling and Covalent Capture

Covalent capture to identify NUAK1 kinase substrates using mass spectrometry was done as previously described (*41*, *67*). Briefly, HA-tagged NUAK1 M132G was purified from HEK293T cells by affinity purification using HA antibody-coated beads. Cells expressing HA-tagged kinase-dead NUAK1-K84M were used as negative controls. Cells were lysed with 25mM HEPES pH 7.2, 150mM KCl, 1x cOmplete protease inhibitor cocktail (Sigma), 1x Halt phosphatase inhibitor (Thermo), 1 mM DTT, 1 mM EDTA, and 1% Triton X-100. Lysate was incubated on ice for 30 min with frequent vortex to lyse cells, followed by centrifugation at 20,000g for 15 min. Supernatant was incubated with 30μL anti-HA resin for 3hr rotating at 4C. 100μl brain lysate at 20mg/ml protein concentration was used for labeling by NUAK1-AS or NUAK1-KD. A total of 5 experimental replicates each for NUAK1-AS or NUAK1-KD were completed with p15 mouse brain lysate. The labeling reaction comprised of 1μM cAMP-dependent protein kinase inhibitor (PKI 5-24, Tocris), 0.2μM protein kinase C inhibitor (Bisindolylmaleimide-I, Calbiochem), 3mM GTP (Sigma), 100μM ATP (Sigma), 1μM okadaic acid (Sigma), 2x Halt Phosphatase inhibitor (ThermoFisher) and 0.5mM Furfuryl-ATP-γ-S (Axxora). Labeling of the brain lysate was quenched by addition of EDTA to 50mM final concentration. The labeled reaction was denatured by adding solid urea (60% w/v) and TCEP (10mM final concentration) and incubating at 55°C for an hour. Samples were then diluted with two volumes of ammonium bicarbonate (Sigma) and two volumes of TCEP (to maintain 10 mM final concentration) and the pH adjusted to 8.0, then digested with Trypsin (Promega, sequencing grade #V5113) with overnight rotation at 37C. Trypsin was quenched the following day by adding 0.5% TFA (Trifluoroacetic acid, Sigma). Peptides were ascertained to be at pH 3.0, then desalted and cleaned using C18 cartridges (Waters Sep-Pak Classic C18 Cartridges), and finally eluted by 70% Acetonitrile (ACN) and 0.1% TFA. Peptide solution was reduced in volume by SpeedVac and final solution adjusted to 50% ACN and 50mM HEPES pH 7.3. Samples were evenly adjusted to pH7.0 using 5% NaOH, incubated with SulfoLink Coupling Resin (ThermoFisher, 20401) and mixed overnight on a nutator at room temperature protected from light. Addition of 25μg Bovine Serum Albumin (BSA) to the beads prior to sample addition ensured low background binding. The following day, the beads were washed over equilibrated fritted columns (Sigma) sequentially using Ultrapure H2O (Thermo), 5M NaCl, 50% ACN, 5% Formic Acid and 1mM DTT. Bound thiophosphopeptides were eluted through hydrolysis reaction by addition of Oxone (1mg/ml Oxone in ddH2O freshly prepared, Sigma 228036). The resulting phosphopeptides were extracted and desalted using C18 StageTips according to the published protocol (*38*, *39*).

#### Reversed-Phase LC-MS/MS Analysis

Peptide samples were separated on an EASY-nLC 1200 System (Thermo Fisher Scientific) using 20 cm long fused silica capillary columns (100 μm ID, laser pulled in-house with Sutter P-2000, Novato CA) packed with 3 μm 120 Å reversed phase C18 beads (Dr. Maisch, Ammerbuch, DE). The LC gradient was 90 min long with 5–35% B at 300 nL/min. LC solvent A was 0.5% (v/v) aq. acetic acid and LC solvent B was 80% acetonitrile in 0.5% (v/v) acetic acid. MS data was collected with a Thermo Fisher Scientific Orbitrap Fusion Lumos using a data-independent acquisition (DIA) method with a 120K resolution Orbitrap MS1 scan and 12 m/z isolation window, 30K resolution Orbitrap HCD MS2 scans for precursors from 400-1000 m/z.

#### Protein and Phosphosite Identification

Data .raw files were converted to .mzML using MSConvert 3.0.24285 and searched using FragPipe version 21.1 and MSFragger version 4.0, with quantification through DIA-NN version 1.8.2 (*68*, *69*). The database search was against the UniProt mouse protein sequence database (downloaded 02-14-2024) with supplemental spike-in of common contaminants, containing 17642 sequences and 17642 reverse-sequence decoys, as well as the sequence for human NUAK1. For the MSFragger analysis, both precursor and initial fragment mass tolerances were set to 20 ppm. Enzyme specificity was set to “stricttrypsin” and up to two missed cleavages were allowed. Oxidized Met, acetylated N-termini, and phosphorylated Ser/Thr/Tyr were set as variable modifications and carbamidomethylated Cys was a fixed modification. The maximum number of variable modifications per peptide was set to 3. FragPipe/DIA-NN output files were processed using a Python script to extract +/-7 AA sequence flanking identified phosphosites from the FASTA file.

#### Database Usage

Developing and adult human brain transcriptomics data were obtained and plotted using HBT database (https://hbatlas.org/). These data were generated by HBT using Affymetrix GeneChip Human Exon 1.0 ST Arrays from over 1340 tissue samples sampled from both hemispheres of postmortem human brains, with specimens ranging from embryonic development to adulthood and representative of both males and females from multiple ethnicities. Allen Brain Atlas was used to search for NUAK1 mRNA expression in the mouse brain (https://mouse.brain-map.org/experiment/show/73497621) as well as PSD3 mRNA expression (https://mouse.brain-map.org/experiment/show/69095967). Evolutionary conservation analysis for both NUAK1 and PSD3 was performed using the ConSurf-DB web server (*32*). Amphipathic helix prediction was performed using the NUAK1 protein sequence from Uniprot and the Heliquest (*35*). NUAK1 consensus sequence and phosphorylation preference for substrates was determined using the publicly available Kinase Library database which was generated using the data from Johnson *et al*, 2023 (*42*).

#### Molecular Modeling of NUAK1 and its Substrates

Prediction of NUAK1(O60285) with PSD3(Q9NYI0) and NUAK1(O60285) with PP1beta(P62140) and MYPT1(O14974) was obtained using Alphafold3 (*33*). Alphafold3 was installed via GitHub (https://github.com/google-deepmind/alphafold3) on a machine running Ubuntu 24.04 LTS with two Nvidia A6000 48GB GPUs. Structural figures were prepared in PyMOL (Schrodinger). Predictions were performed with 20 recycles with 5 diffusion samples per seed. Multiple seeds (217, 275, 428, 551, and 924) were used for random number generation to increase robustness of prediction.

#### Statistics and Reproducibility

Statistical analyses were performed with GraphPad Prism (version 10.0). No statistical method was used to pre-determine sample size, and no data were excluded from the analyses. The experiments were not randomized, and investigators were not blinded during experiments and data analysis. All experiments were performed in at least three independent replicates.

## Supplementary Materials List

Tables S5 to S6

Data files S7

## Acknowledgements

We are grateful for funding support by the National Institutes of Health to SY (R01MH121674, R01MH130336), JS (T32 GM007750) and SEO (R01GM129090, R21CA288806). Funding for the purchase of ELYRA7 Lattice SIM microscope was generously provided by the Murdock Foundation Equipment Grant to SY with matching funds from the University of Washington. This study used an EASY-nLC1200 UHPLC and Thermo Scientific Orbitrap Fusion Lumos Tribrid mass spectrometer purchased with funding from a National Institutes of Health SIG grant S10OD021502 to SEO.

## Author Contributions

J.S. designed and performed all experiments unless otherwise noted below under the supervision of SY. DM ran the proteomics samples on the LC-MS/MS and performed data analysis under the supervision of SEO. SP performed live imaging of PSD3 and PLC-delta-PH in neurons and PSD95 analysis in neurons expressing NUAK1. AS performed dendritic spine morphology and vesicle area quantifications. MC performed Alphafold predictions and PyMol modeling. Manuscript was written by JS and SY and edited by all authors.

## Data and materials availability

All data needed to evaluate the conclusions in the paper are present in the paper or the Supplementary Materials. All materials generated in this study will be shared upon reasonable request to SY.

---

**Figure.S1.**
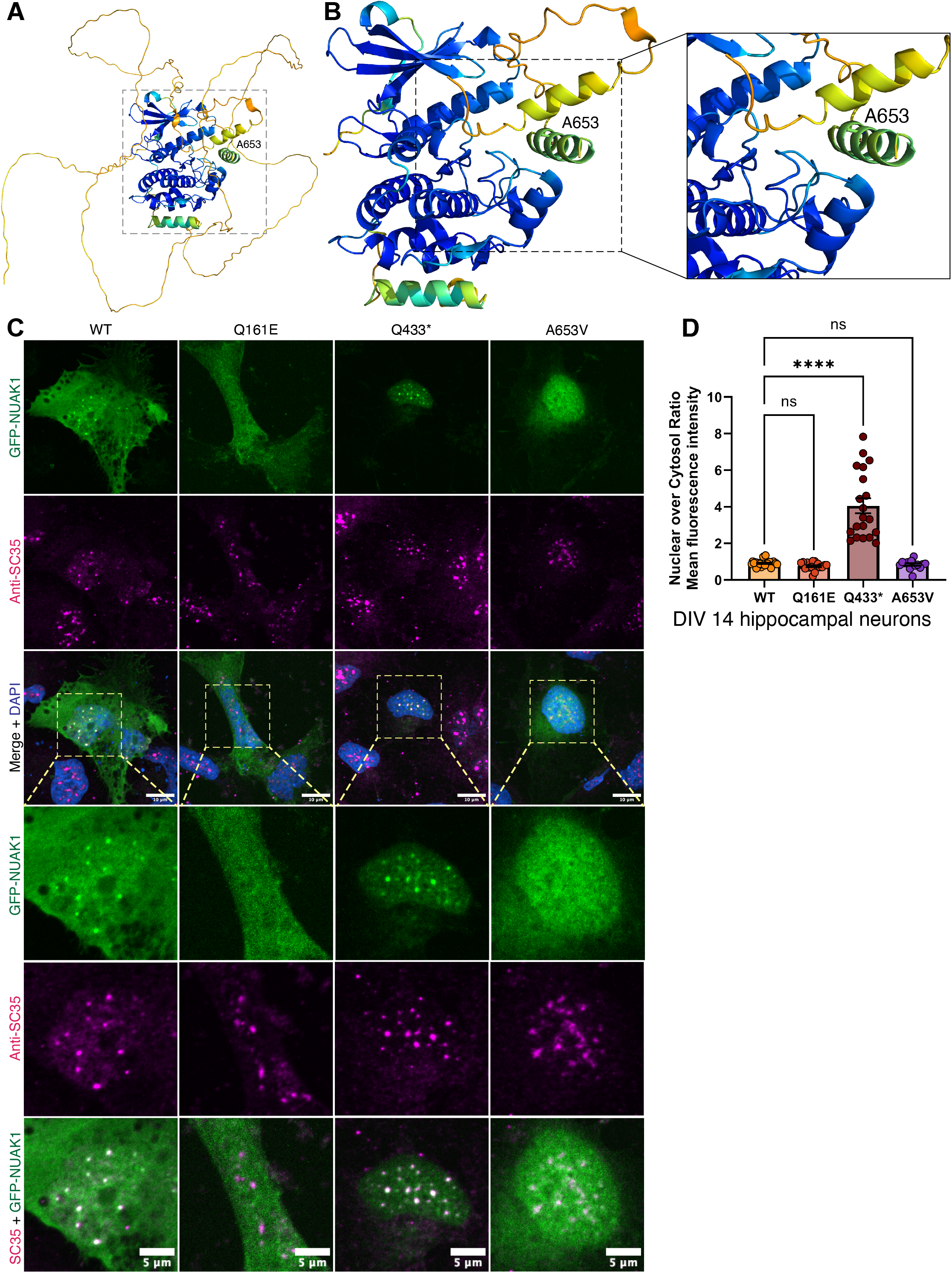

**Figure.S2.**
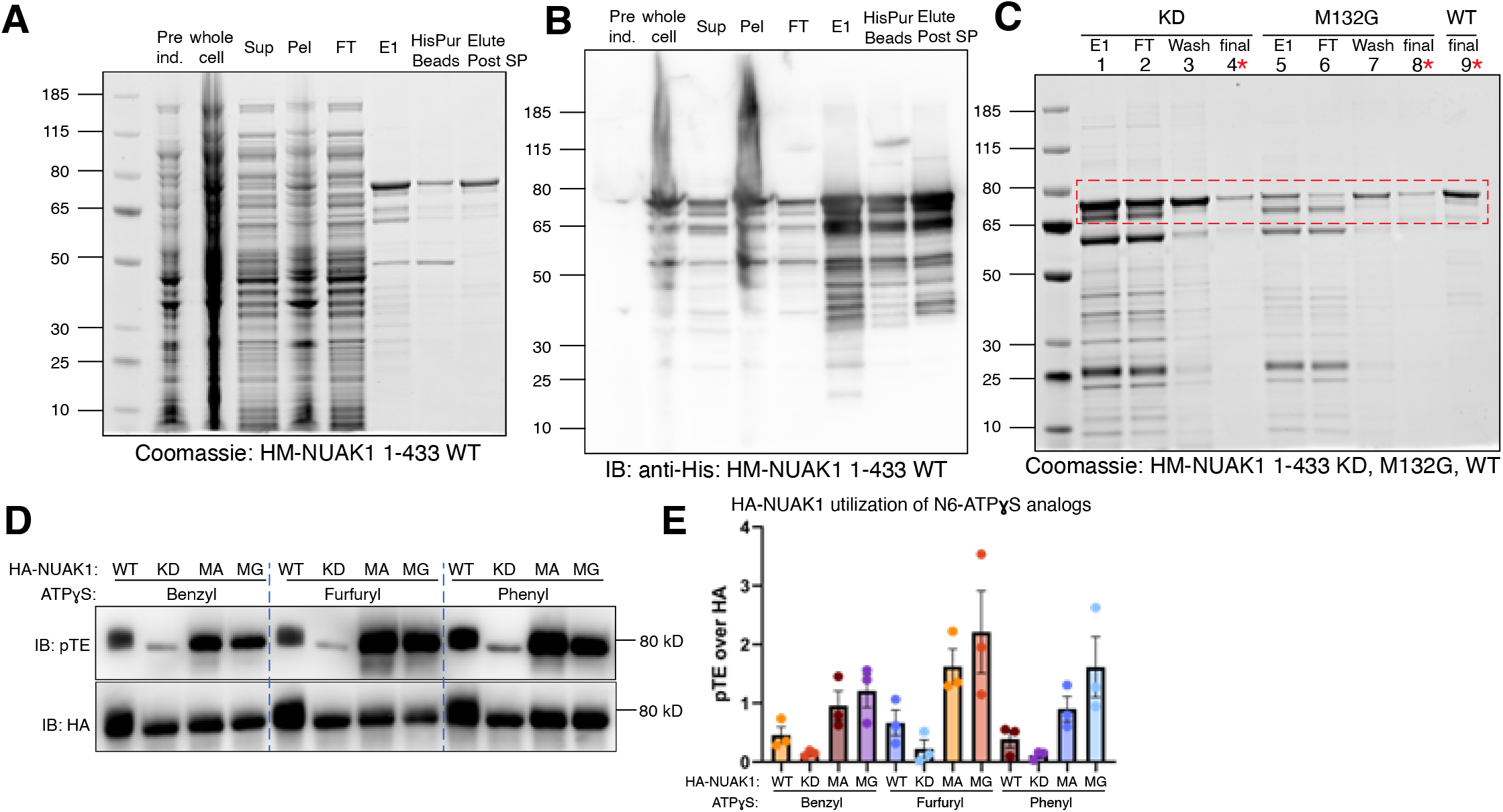

**Figure.S3.**
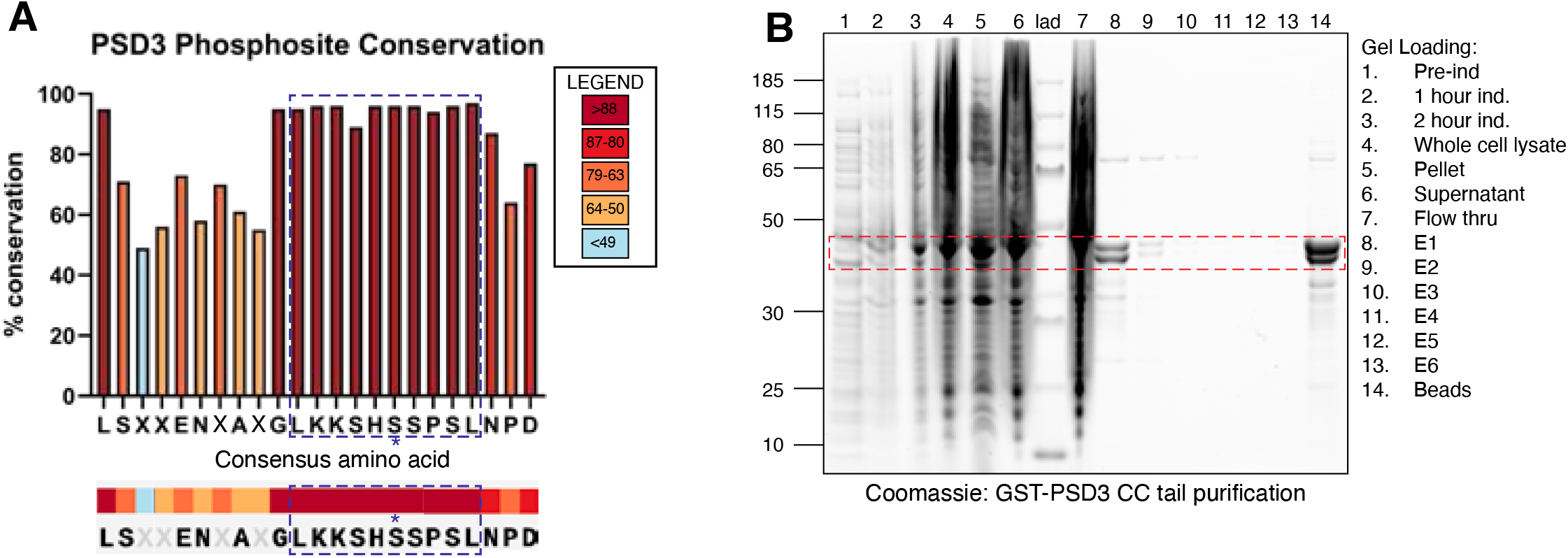

**Figure.S4.**
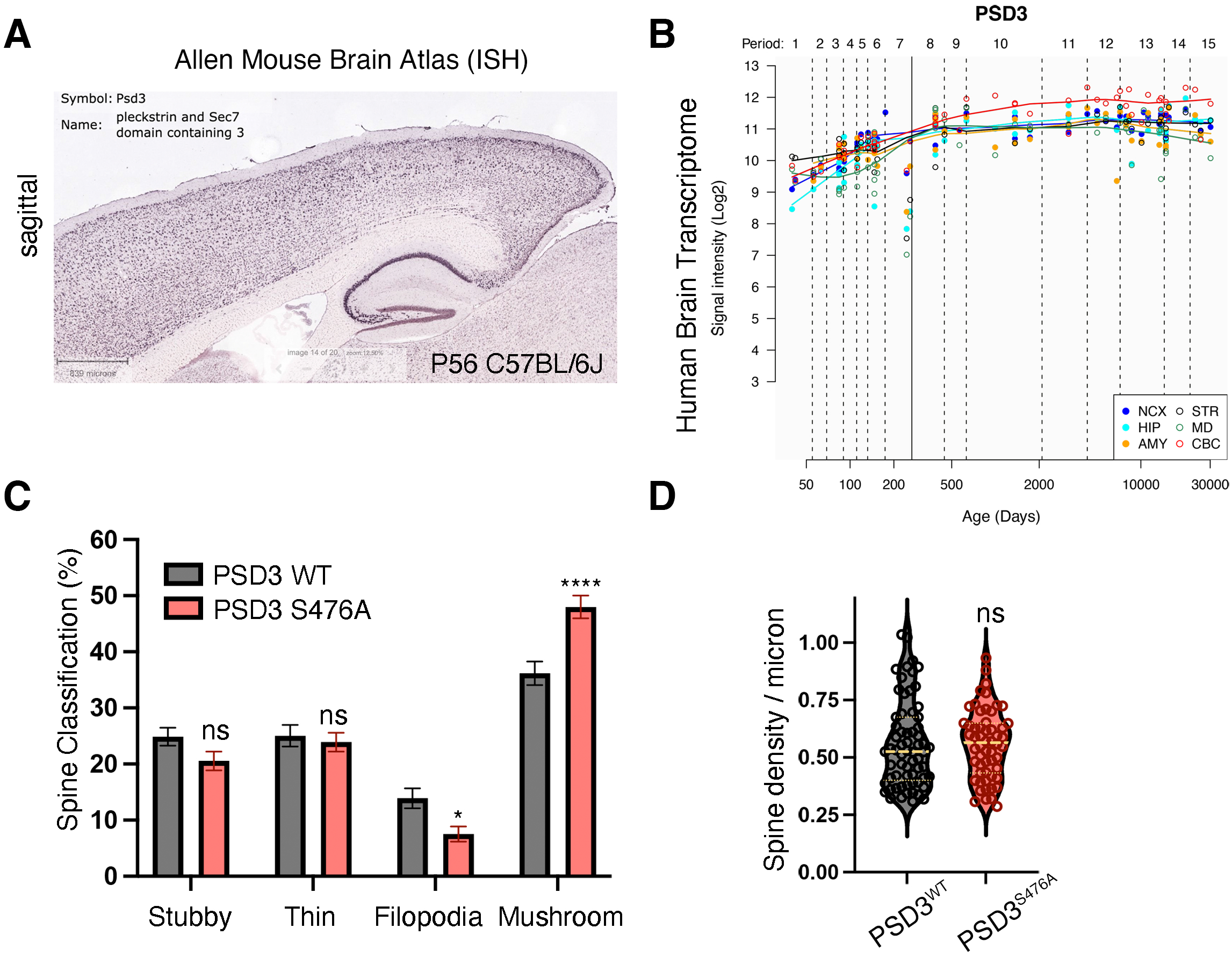

